# Vicarious trauma primes innate immunity and reconfigures human brain networks

**DOI:** 10.64898/2026.01.05.697785

**Authors:** Byeol Kim Lux, Melanie C. Kos, David Ward, Fred W. Kolling, Xin Li, Claudia V. Jakubzick, Tor D. Wager

## Abstract

Vicarious trauma—witnessing serious harm to others—produces systemic neuroimmune effects in animal models. However, its brain, autonomic, and immune consequences in humans remain unclear. We scanned 88 participants with fMRI during a naturalistic movie-viewing paradigm depicting either industrial cruelty to farm animals (Vicarious Trauma, VT) or positive human-animal interactions (Vicarious Community, VC) with concurrent autonomic physiology and gene expression in circulating monocytes. VT, compared to VC, evoked sustained sympathetic arousal lasting 40 minutes post-viewing and primed innate immune cells to elicit elevated pro-inflammatory (upregulating IL1B and CXCL8) and reduced anti-inflammatory (TGFB1) responses. VT reduced integration in a frontoparietal control system and increased coupling between a salience/action-mode system with paralimbic, basal ganglia, and sensorimotor regions. Disrupted negative coupling between frontoparietal/default-mode and hippocampal systems predicted VT-related CXCL8 increases. These findings shed light on how the brain translates vicarious trauma into the priming of innate immunity.

## Introduction

Vicarious trauma—the experience of witnessing trauma or violence to others—is a potent stressor that contributes to moral injury, post-traumatic stress disorder (PTSD), chronic health issues, and reduced prosocial behaviors^1–7^. It is distinct from generic stress and is a common experience in war, caregiving, emergency response, and some occupations. Studies in nonhuman animals have shown that witnessing conspecifics in distress induces hyperalgesia^8^ and acute inflammatory responses^9–12^, persistent depression-like behaviors, social withdrawal^11,13,14^, and increased alcohol and cocaine intake^15,16^. Observing harm to others signals potential danger, which is thought to engage an evolutionarily conserved anticipatory program that coordinates neuroimmune and autonomic responses to prepare for injury and wound repair^17–22^. However, how vicarious trauma in healthy humans induces such anticipatory brain-body states and how these states become biologically embedded remains poorly understood. Here, we used a morally salient, ecologically valid paradigm–a 30 minute documentary showing farm animals being slaughtered in cruel ways, contrasted with a matched film depicting farm animals interacting with humans in playful and loving settings–paired with functional magnetic resonance imaging (fMRI), continuous autonomic monitoring, and peripheral immune cell transcriptomics to reveal linked differences in brain and immune responses.

Despite extensive work on psychosocial stress, few human paradigms have investigated the immersive, morally salient, and personally relevant qualities of real-world vicarious trauma. One line of related research on vicarious trauma has not been experimental, limiting causal inference about underlying physiology^1–5,23^. A second line uses laboratory stressors to modulate inflammatory markers such as interferon-γ, interleukin-1β (IL-1β), tumor necrosis factor-α (TNF-α), and cortisol^24–27^, but these studies are based on general psychosocial stress elicited by negative social evaluation and do not elicit vicarious trauma. A third line models “vicarious distress” with fictional violent movies, injury photographs, or observing others under evaluative threat^28–35^. Although these manipulations can elicit physiological responses, decontextualized or fictional stimuli often lack the authenticity, sustained exposure, and perceived injustice that define vicarious trauma and can blunt empathic engagement^36–38^. In contrast, the moral-emotional aspects of vicarious trauma are particularly associated with moral injury, which is linked to hostility, PTSD, depression, and suicide attempts^6,7,39–41^ and may elicit different immune profiles than other forms of stress. Real-world vicarious trauma evokes guilt and shame^1,42^, and stressors that elicit these emotions produce larger pro-inflammatory responses than otherwise similar stressors^43,44^. These distinctions from other stressors motivate ecologically valid human paradigms that capture morally salient vicarious trauma and its physiological consequences.

To meet this need, we developed a naturalistic film paradigm using two professionally produced 30-minute documentaries assembled from real archival footage. A Vicarious Trauma (VT) film depicts the industrial handling and slaughter of farmed animals common in Western food production, and a Vicarious Community (VC) film showcases the same species’ intelligence, sociality, and positive human-animal interactions. Dynamic, real-world footage evokes stronger emotional and physiological engagement^30,45,46^, particularly when content is perceived as real and personally meaningful^37,47–51^. By tying content to everyday consumption choices, complicity, and preventable suffering, the VT film was designed to elicit empathic engagement, outrage, guilt, and shame, whereas VC reinforced animals’ sentience and prosocial engagement without traumatic elements.

Another critical gap is the lack of integrative, cross-system measurements linking brain, autonomic, and immune responses to vicarious trauma. Given that acute and chronic stress are linked to long-term health outcomes^27,52–54^, understanding the physiological cost of vicarious trauma requires a system-level approach. Nevertheless, most human studies assess brain activity and peripheral physiology in isolation or with limited scope. Although neuroimaging studies on vicarious distress include autonomic measurements such as heart rate or skin conductance, they rarely include immune measures^30,31,33,55,56^. Accordingly, it remains unclear how observing others’ distress engages neural dynamics, autonomic output, and immune programming that may elevate health risk.

Achieving an integrative brain-autonomic-immune understanding of vicarious trauma requires tackling two methodological challenges. The first challenge is measuring brain states that are sustained and dynamic. Simple region-based analyses and brief, trial-based tasks cannot capture the prolonged, fluctuating affect and evolving network interactions that unfold during extended, immersive experience^57–60^. To characterize how brain networks differ between VT and VC, we used template-constrained independent component analysis (ICA; NeuroMark^61–63^) to derive individualized whole-brain network maps from each participant’s 30-minute movie-viewing runs. Seeding with canonical resting-state templates allows us to sensitively capture large-scale systems, while still accommodating interpersonal differences in functional anatomy (a hallmark of the precision functional mapping approach^64^). From these participant-specific components, we estimated both stable and time-varying coupling among large-scale networks such as frontoparietal control, default-mode, and salience/action-mode networks, enabling comparison of how these networks reconfigured between VT and VC.

The second challenge is measuring immune responses with sufficient cellular specificity to be meaningfully integrated with fMRI. Most psychosocial stress research measures circulating hormones or cytokines (for example, serum or salivary cortisol)^26,65–69^. However, these mediators are produced by multiple cell types and play multiple, overlapping roles in innate and adaptive immunity^70–72^, so circulating levels provide limited insight into which immune populations are being tuned^73^. To overcome this, we performed RNA sequencing on enriched circulating monocytes collected before and after each viewing session, along with fMRI scans and autonomic recordings. Monocytes are first responder cells in the innate immune system^74^. They are rapidly mobilized by stress signals, infiltrate tissues, and initiate pro-inflammatory cascades (IL-1β, IL-8, and TNF-α) and reparative signaling (TGF-β1), including neutrophil recruitment to sites of injury via the chemokine IL-8^75–78^. Monocyte transcriptional state provides a sensitive readout of early immune modulation that can precede detectable changes in circulating proteins^79–81^. Although T cells can also secrete some of these factors, their activation is antigen-specific and thus reflects adaptive rather than innate immunity^82^. Thus, monocytes are particularly well-positioned to show anticipatory responses to vicarious trauma, guided by internal representations of injury and threat.

By linking neural network indices and autonomic metrics to monocyte gene expression changes, we tested whether specific patterns of large-scale brain coupling predict peripheral immune responses to vicarious trauma. We hypothesized that morally salient vicarious trauma would evoke anticipatory shifts in monocyte programming, even in the absence of physical injury, biasing the system toward a pro-inflammatory state of readiness. Guided by prior work in stress immunology^26,69,83–85^, we selected an a priori panel of cytokines and chemokines prominently expressed by monocytes and robustly detected in our samples: IL1B, CXCL8, and TNF (pro-inflammatory), and TGFB1 (anti-inflammatory). We quantified within-subject transcriptional changes in these targets and examined their coupling with heart-rate fluctuations and large-scale network reconfiguration to reveal coordinated neuroimmune responses to vicarious trauma.

Our multimodal results support this account. VT induced persistent sympathetic activation, with elevated heart rate persisting for at least 40 minutes after movie viewing. Immunologically, monocyte RNA-seq revealed a proinflammatory shift under VT relative to VC–upregulation of IL1B and CXCL8 and downregulation of TGFB1, with no significant change in TNF–consistent with selective, anticipatory innate immune programming. Neurally, VT shifted large-scale organization from a state more coordinated by executive-control systems to one more driven by visceral and motivational circuits: cohesion within Language-Control systems and their coupling to Memory-default and Affective-Action systems weakened, while connections tightened between Affective-Action system and interoceptive and basal ganglia pathways. Critically, a specific connectivity pattern integrating visual, default-mode, paralimbic, and insular/temporal nodes predicted VT-related increases in CXCL8, linking how they saw and internalized the trauma to downstream innate immune responses. Together, these findings demonstrate integrative engagement of brain, autonomic, and immune systems during ecologically valid vicarious trauma and identify paralimbic network reconfiguration as a neural correlate of monocyte inflammatory programming.

## Results

### Study Design and Affective Ratings

The fMRI experiment consisted of two sessions (mean interval = 9.88 ± 9.48 days). Each session began with a peripheral blood draw for baseline measures, followed by fMRI scanning that included three movie-viewing task runs, a 10-minute resting-state run, and additional tasks not analyzed here (**Fig. 1A**). Heart rate and skin conductance were recorded continuously during scanning. After the scan, participants completed a 5-minute resting autonomic recording of heart rate and skin conductance in a behavioral testing room. A second blood sample was then taken for bulk RNA sequencing of isolated monocytes. This multimodal dataset enables a systems-level and mechanistic evaluation of Vicarious Trauma (VT) versus Vicarious Community (VC) across neural, autonomic, and immune/transcriptomic measures.

**Figure 1.**
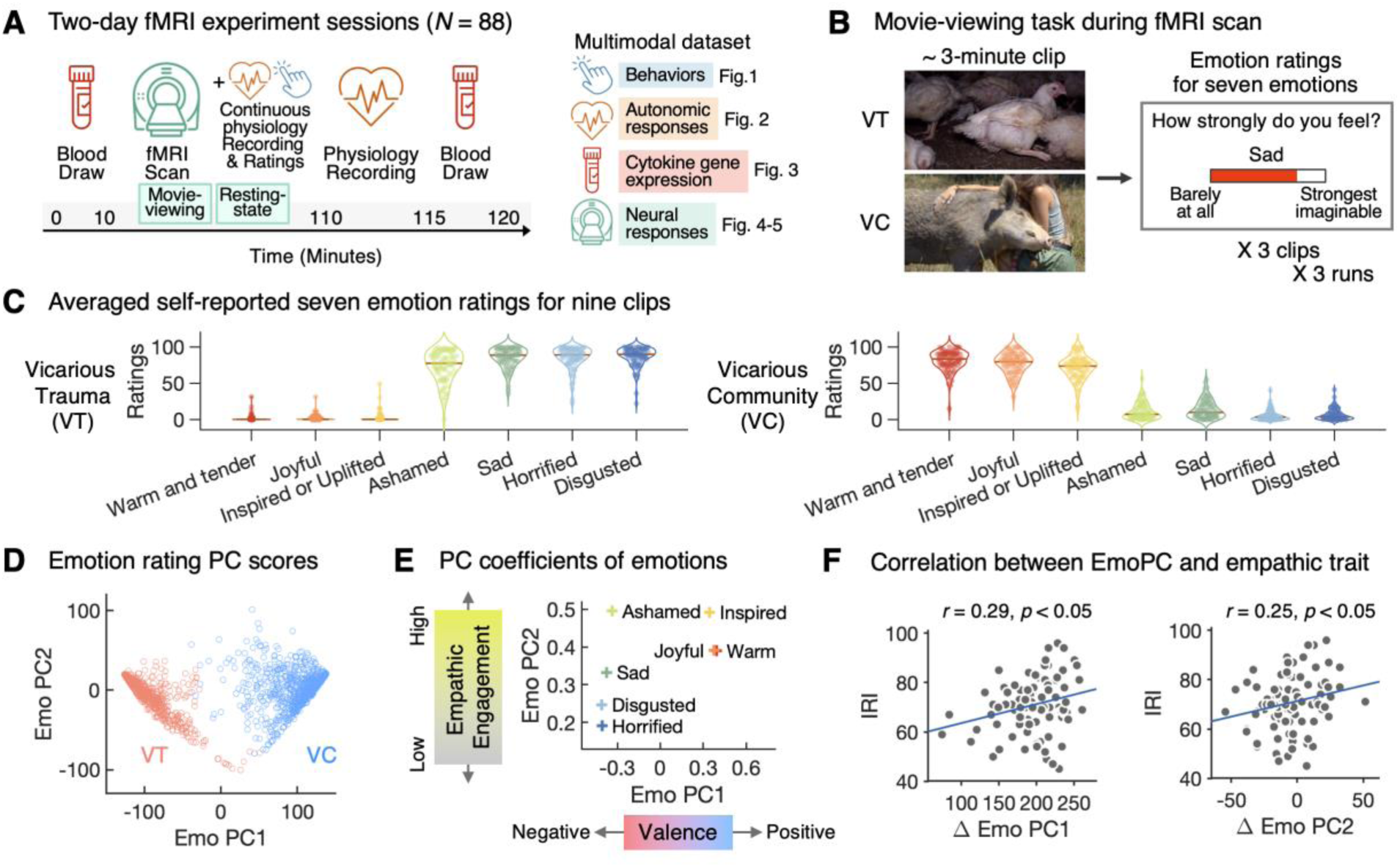
Experimental overview and self-report emotion ratings. (**A**) Participants completed a two-session fMRI experiment conducted on two different days, including two peripheral blood draws (pre- and post-scan), fMRI scanning (three movie-viewing runs and a 10-minute resting-state run), and a 5-minute post-scan resting physiological recording of heart rate and skin conductance. During scanning, heart rate and skin conductance were recorded continuously, and self-reported emotion ratings were collected intermittently. The time bar indicates the time elapsed since the first blood draw. This design enables a multimodal assessment of psychoneuroimmunological responses to VT versus VC. (**B**) In each session, participants watched one 30-minute animal documentary (VT or VC). The film was divided into nine ∼3-minute clips. After each clip, they rated seven emotions: “warm and tender,” “joyful,” “inspired or uplifted,” “ashamed,” “sad,” “horrified,” and “disgusted.” (**C**) VT and VC elicited opposing emotion profiles. VT evoked higher ratings of negative emotions (“ashamed,” “sad,” “horrified,” “disgusted”), whereas VC evoked higher levels of positive emotions (“warm and tender,” “joyful”) and higher levels of “inspired or uplifted.” (**D**) Principal component analysis of clip-level ratings across participants revealed two principal components. The first component (PC1) separated VT from VC. (**E**) Emotion loadings indicated that PC1 captured valence (higher scores = more positive valence). PC2 loaded highest on “ashamed” and “inspired or uplifted” and lowest on “horrified” and “disgusted,” suggesting an empathic engagement dimension; we therefore refer to them as EmoPC1 (valence) and EmoPC2 (empathic engagement). (**F**) Condition differences in EmoPC1 and EmoPC2 (VT−VC) correlated with the total score of the Interpersonal Reactivity Index (IRI), such that higher trait empathy was associated with larger VT−VC differences in both EmoPCs.

During the movie-viewing task, participants watched one of the two 30-minute documentary conditions (VT or VC), counterbalanced across participants (**Fig. 1B**). Each documentary film was segmented into nine clips of approximately three minutes. After each clip, participants rated their emotional responses to each clip across seven emotions: “warm and tender,” “joyful,” “inspired or uplifted,” “ashamed,” “sad,” “horrified,” and “disgusted”. These dimensions were selected to capture basic and morally relevant emotions implicated in VT and VC. As expected, the VT videos elicited high levels of negative emotions (“ashamed,” “sad,” “horrified,” and “disgusted”) and low levels of positive emotions (“warm and tender” and “joyful”) and low “inspired or uplifted” feelings (**Fig. 1C**, **fig. S1**). In contrast, the VC videos robustly evoked high levels of positive emotions and “inspired or uplifted” feelings, and low levels of negative emotions (**table S1**).

Principal component analysis was performed on the emotion ratings across all clips and participants to identify key dimensions of affective responses (**Fig. 1D**). The first principal component (EmoPC1) represented emotional valence, clearly separating ratings from the VT and VC (explained 89.79% of the variance). Positive emotions loaded positively, while negative emotions loaded negatively (**Fig. 1E**). The variables that contributed the most to the second principal component (explained 89.79% of the variance) were “inspired or uplifted” and “ashamed,” suggesting that EmoPC2 reflects empathetic engagement, while “horrified” and “disgusted” showed the lowest loadings. Participants had nine EmoPC scores for each condition, and these scores were significantly correlated across conditions (**fig. S1**; *Ps <* 0.01). Their VT−VC difference scores (ΔEmoPC) were also significantly correlated with EmoPC scores in VT (**Fig. S1**; *Ps <* 0.001), suggesting that the difference scores capture participants’ reactivity to the movies. For EmoPC1, individuals who showed stronger negative valence responses during VT tended to show more positive responses during VC, resulting in larger VT−VC contrasts (ΔEmoPC1). For EmoPC2, individuals with greater empathic engagement during VT also tended to show greater engagement during VC and larger VT−VC differences (ΔEmoPC2). Moreover, ΔEmoPC values were significantly correlated with the total score of the Interpersonal Reactivity Index (IRI) (*Ps <* 0.05), suggesting that individuals with greater empathic traits showed larger differences in both EmoPCs across conditions (**Fig. 1F**).

### Persistent Sympathetic Activation in VT vs. VC

Relative to VC, VT produced stronger sympathetic activation, consistent with elevated stress-related autonomic responses. Heart rate, measured via electrocardiogram (ECG), was higher during VT than during VC (**Fig. 2A-B**; paired *t*-test, *t* = 2.40, *P* = 0.019, *n* = 85). This heart rate elevation persisted beyond the movie runs, remaining higher during the subsequent 10-minute resting-state fMRI run (**Fig. 2B**; **table S2**; *t* = 2.98, *P* = 0.004) as well as during the post-scan 5-minute resting physiology recordings approximately 40 minutes after the movies ended (*t* = 2.41, *P* = 0.018). During the brief pre-movie baseline (−20 to 0 s), heart rate was slightly higher in VT than VC, but the difference was not significant (*P* > 0.05). Baseline was measured immediately before movie onset, after participants had been informed of the upcoming condition and the scanner had started. We investigated whether self-reported emotions, summarized by EmoPCs, predicted heart rate during movie watching using a linear mixed-effects model. The model revealed a significant interaction between EmoPC1 and EmoPC2 in predicting heart rate (**Fig. 2C**, **table S3**; *β* = −0.00029, *t* = −3.36, *P* = 0.001, *n* = 85). Empathic engagement (EmoPC2) moderated the association between valence (EmoPC1) and heart rate during movie-viewing. When empathic engagement was high, more negative valence was associated with higher heart rate; when empathic engagement was low, valence was only weakly related to heart rate.

**Figure 2.**
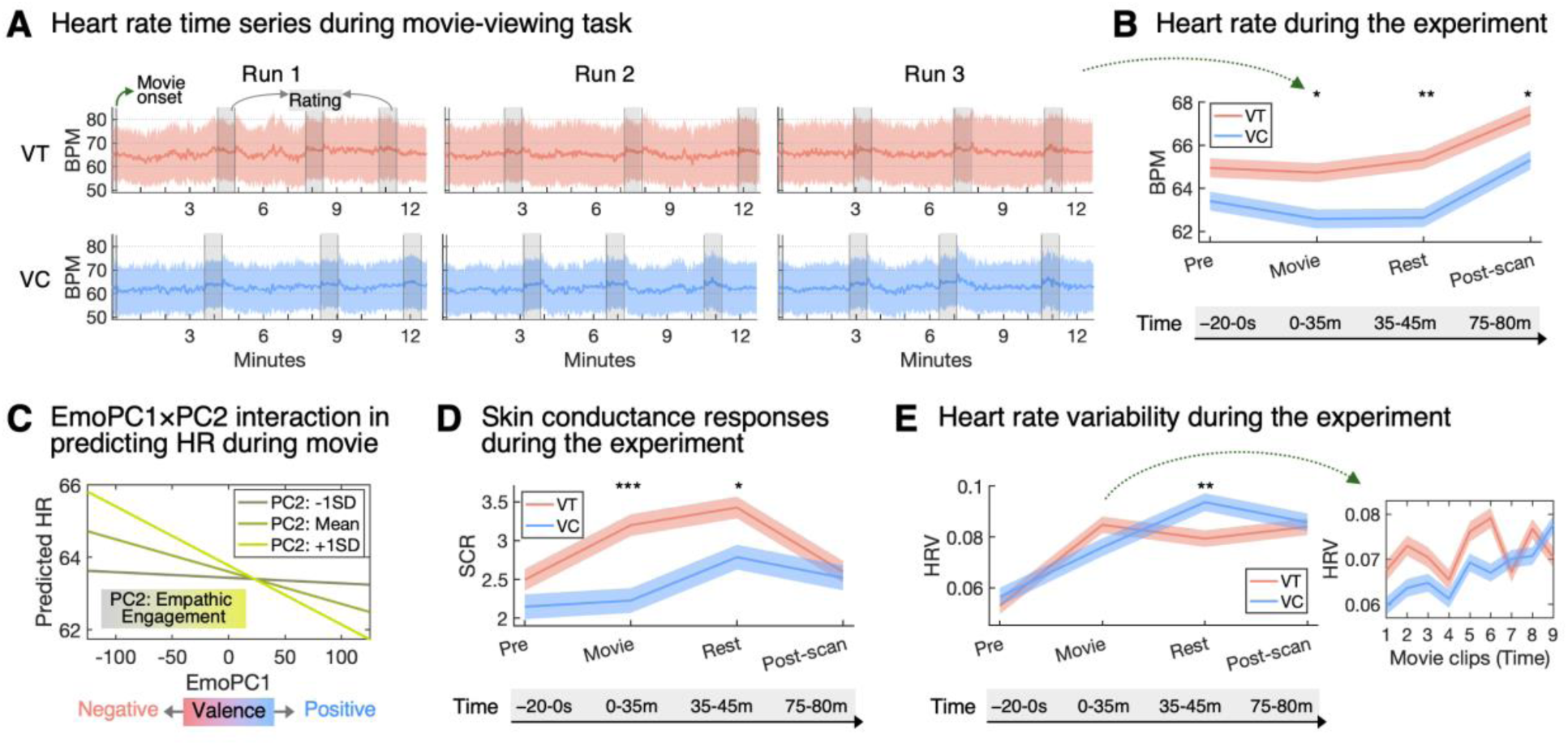
Autonomic responses during Vicarious Trauma and Vicarious Community. (**A**) Heart rate (beats per minute; BPM) was recorded continuously during the in-scanner movie-viewing task (*n* = 85), which consisted of three 12-minute runs. Each run consisted of 3 of the 9 total VT or VC clips. Gray shaded boxes indicate emotion rating periods (three per run). Shaded bands indicate the standard deviation across participants. The y-axis range is identical across conditions, illustrating an overall higher heart rate in VT than in VC. (**B**) Heart rate was compared between conditions across time points: a 20-s pre-movie baseline (Pre), the movie-viewing period (Movie), a 10-minute resting-state run following the movie task in the scanner (Rest), and a 5-minute post-scan resting recording in the behavioral room (Post-scan). Paired *t*-tests revealed a higher heart rate in VT than in VC during the movie, the resting-state run, and even the post-scan recording. The time bar indicates the time elapsed since the movie onset (**table S2**). Lines show the condition means. Shaded bands indicate within-subject variability, consistent with other panels (Fig. 2D**-E**). (**C**) Empathetic engagement (EmoPC2) moderated the association between valence (EmoPC1) and heart rate during movie-viewing. When participants reported high empathic engagement with a clip, more negative valence was associated with higher heart rate. The y-axis shows predicted heart rate, and the x-axis shows EmoPC1. The middle green line depicts the predicted heart rate-EmoPC1 relationship at EmoPC2 = mean. Bright and dark green lines depict EmoPC2 = mean ± 1 standard deviation (SD). (**D**) Skin conductance responses (SCR) were measured over the same timeline and compared using paired *t*-tests (*n* = 72). SCR was greater in VT than in VC during movie-viewing and the subsequent resting-state run. (**E**) Heart rate variability (HRV) was computed across the experiment. HRV was lower in VT than in VC during the resting-state run but not at other time points (left). The right plot shows HRV across nine movie clips. HRV during VC tended to increase over time, whereas HRV during VT fluctuated across clips. Asterisks denote significance: \**P* < 0.05, \*\**P* < 0.01, \*\*\**P* < 0.001.

Sympathetic activation with VT was also evident in skin conductance and heart rate variability. Skin conductance response (SCR), measured as electrodermal activity (EDA), was higher during VT (**Fig. 2D**, **table S2**; paired *t*-test, *t* = 3.95, *P* < 0.001, *n* = 72) and during the following resting-state run (*t* = 2.51, *P* = 0.015), although no significant differences were observed in the post-scan recording. Although heart rate variability (HRV, measured as the standard deviation of NN intervals) did not differ significantly during movie-viewing, it was significantly lower during the resting-state run after VT compared with VC (**Fig. 2E**; paired *t*-test, *t* = −2.73, *P* = 0.008, *n* = 85). Throughout the movie-viewing task, HRV fluctuated in the VT condition but tended to increase steadily over time in the VC condition. Given that HRV was lower at the pre-movie baseline than during the movie-viewing period, this fluctuation under VT implies disrupted HRV recovery within the in-scanner context. These findings suggest a persistent autonomic threat response in VT and show that empathic engagement modulates movie-evoked sympathetic activation.

### Targeted analysis of stress-responsive cytokines

To examine monocyte gene expression, we enriched monocytes from peripheral blood and analyzed both unenriched and monocyte-enriched samples by flow cytometry (*n* = 55). After gating on live cells, subsets were identified using CD14 versus CD16. SSC versus CD16 distinguished high-SSC CD16^high^ neutrophils, intermediate SSC monocytes, and low-SSC lymphocytes, containing low-SSC CD16^+^ NK cells. Enriched samples showed a marked increase in CD14^+^ monocytes compared to unenriched samples. Thus, the enrichment kit was highly effective at isolating CD14^+^ monocytes (**Fig. 3A**; **fig. S2**). Orthogonal RNA-seq validation (**Fig. 3B**) showed TPM values for canonical lineage markers: high expression of CD14 and negligible expression for CD3E (T cells), CD19 (B cells), MPO (neutrophils), NKG7 (NK cells), and CLEC9A (dendritic cells), consistent with minimal contamination in the monocyte-enriched fraction. Following confirmation of enrichment, we performed bulk RNA-sequencing on the enriched samples.

**Figure 3.**
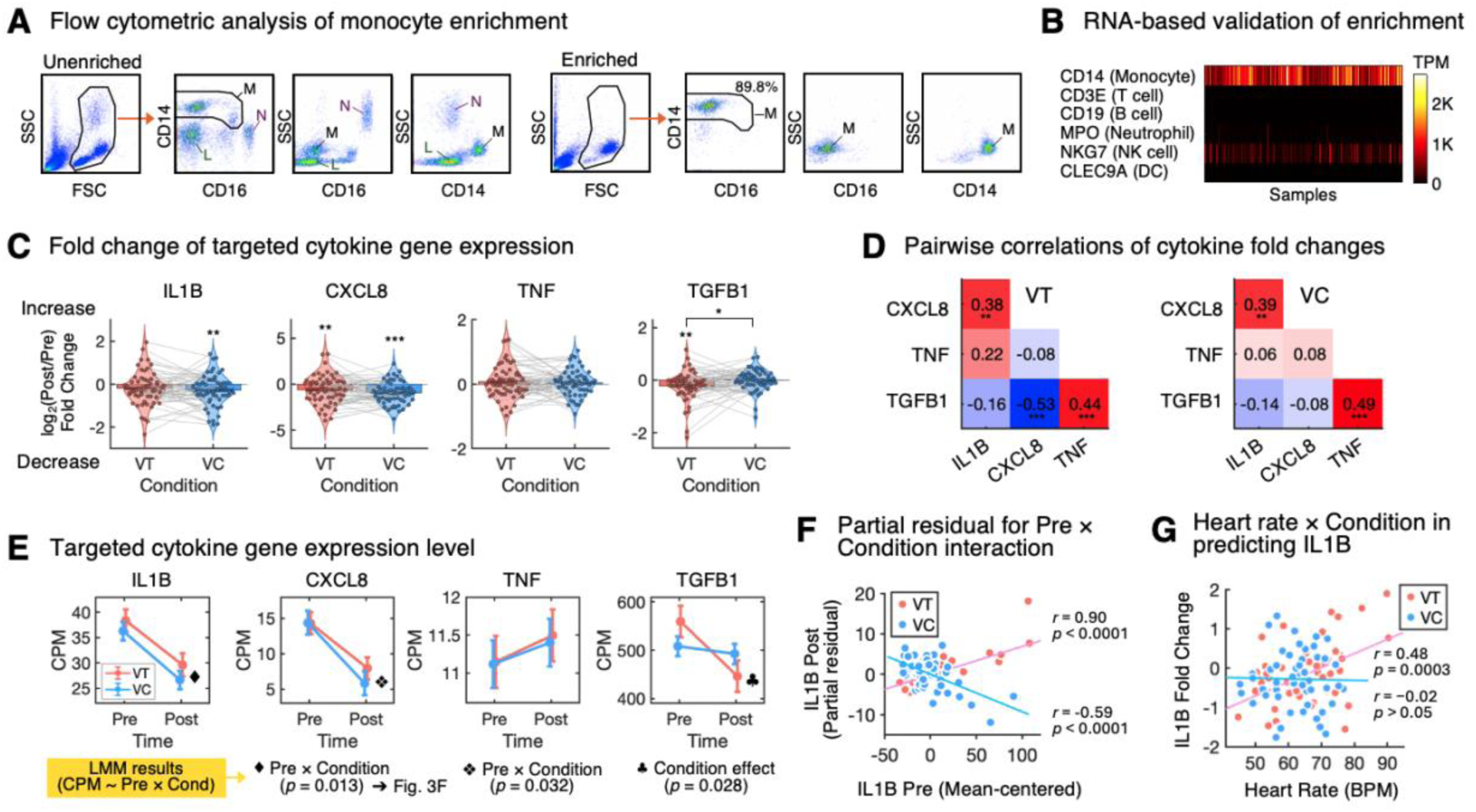
Monocyte-focused RNA-seq identifies VT-related IL1B/CXCL8 upregulation and TGFB1 suppression. (**A**) Representative flow cytometry plots illustrating monocyte enrichment. The left four panels show an unenriched sample, and the right four panels show the corresponding enriched sample. After excluding dead cells and doublets, the remaining live cells were plotted in FSC vs SSC to gate live cells (black outline). Gated live cells were then plotted as CD14 vs CD16 and as SSC vs CD16 or CD14. CD14 vs CD16 illustrates classical (CD14++CD16−), intermediate (CD14++CD16+), and non-classical (CD14+CD16++) monocyte subsets (M, black outline) along with CD16^high^ neutrophils (N) and CD14^−^ lymphocytes (L). SSC vs CD16 and SSC vs CD14 plots distinguish monocytes (intermediate SSC) from high-SSC granulocytes and low-SSC lymphocytes. Enrichment increased the monocyte gate from 24.6 to 89.8%, confirming successful monocyte isolation. (**B**) Orthogonal RNA-seq validation using TPM-normalized expression of lineage markers. The heatmap shows consistently high CD14 expression across samples with negligible expression of CD3E (T cells), CD19 (B cells), MPO (neutrophils), NKG7 (NK cells), and CLEC9A (dendritic cells), supporting effective monocyte enrichment with minimal contamination. (**C**) Violin plots of fold change (FC) for IL1B, CXCL8, TNF, and TGFB1 by condition. Gray lines connect individuals. One-sample *t*-tests on the FC showed significant reductions for IL1B in VC only, CXCL8 in both conditions, and TGFB1 in VT only. Paired *t*-tests identified TGFB1 as the only cytokine with a significant VT−VC difference (VT < VC). (**D**) Correlation matrices showing covariation of cytokine fold changes by condition (numbers are the Pearson *r*). IL1B and CXCL8 were positively correlated in both conditions. CXCL8 and TGFB1 were negatively correlated in VT only. TGFB1 and TNF were positively correlated in both conditions. (**E**) Averaged gene expression (CPM) across time for each cytokine by condition. Error bars indicate within-subject variability. Linear mixed-effects models predicting post-scan expression level from baseline level (Pre) and condition, adjusting for covariates, revealed significant Pre × Condition interactions for IL1B and CXCL8, and a significant main effect of condition for TGFB1. (**F**) Visualization of the Pre × Condition interaction for IL1B: Partial residuals of post-scan expression after removing the main effects of baseline and covariates. Baseline expression strongly predicted higher post-scan levels in VT (*r* = 0.90, *P* < 0.0001) and lower levels in VC (*r* = −0.59, *P* < 0.0001). Each point indicates a participant for each condition, and lines show fitted regressions. (**G**) IL1B fold change correlated with heart rate during movie viewing in VT but not in VC, indicating that the heart rates-IL1B association was specific to VT. Asterisks denote significance: \**P* < 0.05, \*\**P* < 0.01, \*\*\**P* < 0.001

With the bulk RNA-sequencing on the enriched monocytes (expression quantified as counts per million, CPM), we assessed whether gene expression changes in four cytokines (IL1B, CXCL8, TNF, TGFB1) differed between VT and VC. Baseline (Pre-viewing) expression levels of IL1B, TNF, and TGFB1 were stable within-person across conditions (**table S4**; *r* = 0.274-0.355, *Ps* < 0.05), whereas CXCL8 did not, with a weaker correlation (*r* = 0.253, *P* = 0.063). First, one-sample *t*-tests on fold change values (log_2_(Post/Pre); **Fig. 3C**) showed a significant reduction in IL1B after VC (*m* = −0.284, *t* = −2.861, *P* = 0.006), but not after the VT. CXCL8 decreased significantly after both VT (*m* = −0.604, *t* = −2.824, *P* = 0.007) and VC (*m* = −0.957, *t* = −5.790, *P* < 0.001). TNF showed no significant changes in either condition. TGFB1 was significantly reduced after VT (*m* = −0.212, *t* = −2.792, *P* = 0.007), and the VT−VC difference was also significant in a paired *t*-test (*m* = −0.213, *t* = −2.502, *P* = 0.015, see **table S5** for full results). Although IL1B and CXCL8 declined more after VT than VC in one-sample tests, only TGFB1 showed a significant within-subject difference between conditions, reflecting lower post-VT expression.

To assess whether cytokine fold changes covaried, we computed correlations among cytokine fold changes within each condition (**Fig. 3D**). CXCL8 and IL1B were positively correlated in both VT (*r* = 0.382, *P* = 0.004) and VC (*r* = 0.385, *P* = 0.037), consistent with a condition-invariant pro-inflammatory coupling. In contrast, TGFB1 and CXCL8 were negatively correlated in the VT (*r* = −0.530, *P* < 0.001), but not in the VC (*r* = −0.077, *P* > 0.05), suggesting a VT-specific anti-inflammatory axis. TNF and TGFB1 were positively correlated in both VT (*r* = 0.438, *P* < 0.001) and VC (*r* = 0.489, *P* < 0.001).

To model repeated-measure structure and between-subject heterogeneity in condition effects, we fit linear mixed-effects models predicting post-scan gene expression (Post) from baseline expression (Pre) and condition, controlling for randomized condition order, sex, age, and batch (**Fig. 3E**, **table S6** for full results). Baseline expression robustly predicted post-scan levels for all four cytokines (all *β*s > 0, all *P*s < 0.0001), indicating strong within-person stability. IL1B and CXCL8 showed no main effect of condition, but showed significant condition × baseline interactions (*P* = 0.013 and 0.032, respectively). Partial residual for these condition × baseline interactions (controlling for baseline and covariates; **Fig. 3F**) revealed a crossover pattern: higher baseline predicted higher post-scan expression after VT but lower expression after VC, for both IL1B and CXCL8. For TGFB1, a significant main effect of condition suggested lower expression after VT than after VC (*P* < 0.05; **Fig. 3E**), controlling for baseline, with no significant condition × baseline interaction.

Additionally, we examined whether subjective emotional experience (EmoPCs) and autonomic state (heart rate) during movie viewing predicted cytokine change. Mixed-effects models of fold change, including EmoPC1 (valence), EmoPC2 (empathic engagement), and heart rate (controlling for condition order, sex, and age; see **table S7** for full results), revealed that heart rate significantly predicted IL1B fold change (*β* = 0.024, *P* = 0.006). EmoPC1 significantly predicted TGFB1 fold change (*β* = 0.001, *P* = 0.017), consistent with the VT-related reduction. No significant effects were found for CXCL8 or TNF. In a model adding interactions for IL1B (**table S8**), both the main effect of heart rate (*β* = 0.019, *P* = 0.032) and the EmoPC1 × heart rate interaction were significant (*P* = 0.047; **Fig. 3E**), indicating that the positive association between heart rate and IL1B fold change was stronger at more negative valence. Because EmoPC1 closely tracks condition at the session level, this pattern suggests that the heart rate-IL1B association was specific to VT, rather than VC.

### Functional Network Connectivity during movie-viewing

By applying template-constrained ICA (NeuroMark) on our movie-viewing fMRI data, we estimated participant-specific time courses and spatial maps for 105 intrinsic connectivity networks (ICNs; **fig. S3**). Participants’ ICN spatial maps were highly stable across runs in two conditions (**table S9**; mean within-condition spatial correlation: Fisher *z* = 0.756; cross-condition: Fisher *z* = 0.732). For each participant and condition, we computed static functional network connectivity (sFNC; Fisher *z*-transformed correlations between ICN pairs) and contrasted VT with VC using paired *t*-tests (**Fig. 4A**, **fig. S4**). VT, compared to VC, produced widespread significant changes in connectivity: 659 of 5,460 unique ICN pairs survived Bonferroni correction (*q* < 0.05; **Fig. 4B**, **fig. S4**).

**Figure 4.**
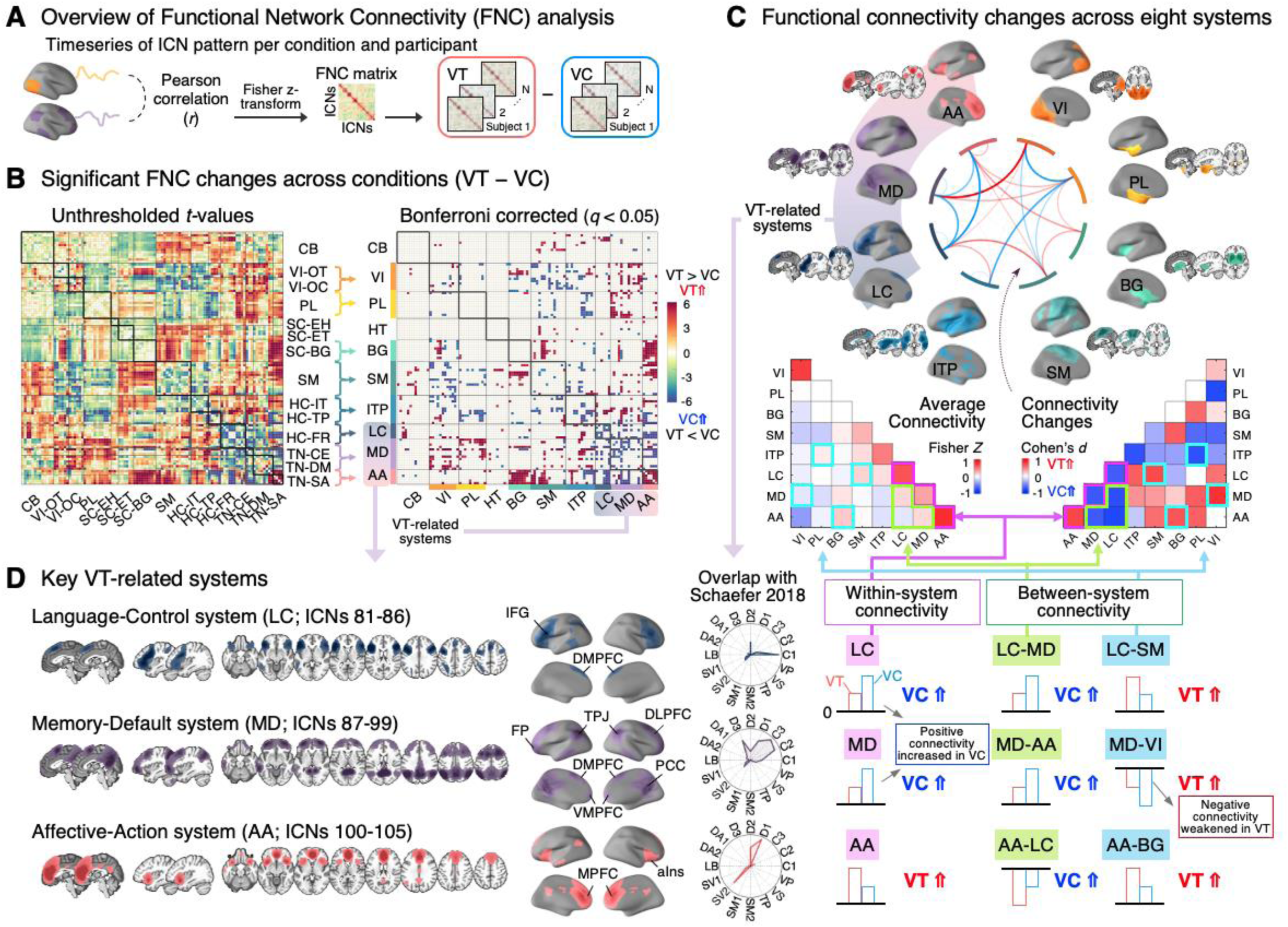
Vicarious Trauma reorganizes large-scale functional network connectivity. (**A**) Using the NeuroMark approach, we estimated participant-level ICN spatial maps and time courses from movie-viewing fMRI. Static FNC was computed as Fisher *z*-transformed pairwise Pearson correlations among ICN time courses within each condition. Condition differences (VT−VC) were tested with within-subject paired *t*-tests across 5,460 pairs. (**B**) Pairwise results (red: VT > VC; blue: VC > VT). Left: *t*-value matrix organized by the 14 NeuroMark subdomains (full list in **fig. S3**). Right: Bonferroni-significant results (*P* < 0.05). This Bonferroni-corrected *t*-value matrix highlights 659 significant pairs (*P* < 0.05; ∣*t*∣ > 4.71). For systems-level interpretation, ICNs were regrouped into the eight systems: Visual (VI), Paralimbic (PL), Basal Ganglia (BG), Sensorimotor (SM), Insular-Temporoparietal (ITP), Language-Control (LC), Memory-Default integration (MD), and Affective-Action integration (AA) (more details in **figs. S4-6**). CB, Cerebellar; HT, Hippocampal-Thalamic. (**C**) Circos diagram showing directions of connectivity changes across the eight systems (red: VT > VC; blue: VC > VT). Brain maps show the spatial distribution of each system (darker color indicates greater overlap across ICNs). Lower-left matrix: mean connectivity among systems (mean of VT and VC; Fisher *z*). Lower-right matrix: Cohen’s *d* for VT−VC connectivity differences (**table S10**). Key findings: LC and MD showed greater within-system connectivity in VC, but AA showed greater within-system coherence in VT. The largest VT > VC increases were observed for MD-VI, AA-BG, and within-AA; the largest VT < VC decreases were observed for VI-PL, ITP-PL, LC-MD, and LC-AA. VT also showed enhanced coupling of AA with PL, BG, SM, and ITP, whereas VC showed stronger within-system connectivity in ITP, LC, and MD. (**D**) Three VT-related systems (LC, MD, and AA). Brain region abbreviations: IFG, inferior frontal gyrus; DMPFC, dorsomedial prefrontal cortex; FP, frontal pole; TPJ, temporoparietal junction; DLPFC, dorsolateral prefrontal cortex; PCC, posterior cingulate cortex; VMPFC, ventromedial prefrontal cortex; aIns, anterior insular cortex; MPFC, medial prefrontal cortex. Radial plots show percent overlap with the 17-network parcellation from^87^. Large-scale network abbreviations: C1-3, Control A-C; D1-3, Default A-C; DA1-2, Dorsal Attention A-B; LB, Limbic; SV1-2, Salience/Ventral Attention A-B; SM1-2, Somatomotor A-B. Right panels show schematic bars summarizing connectivity by condition, indicating whether connectivity is positive or negative and greater in VT or VC (red bar = VT, blue bar = VC, black line = connectivity 0). VT reduced coupling among VT-related systems and within-LC/MD, while increasing within-AA and selective between-system coupling (e.g., MD-VI, AA-BG).

To enable systems-level interpretation of these task-evoked reconfigurations, we regrouped the ICNs into a smaller set of interpretable systems. Since the NeuroMark template was derived from resting-state data, we did not expect a one-to-one mapping onto networks engaged during naturalistic movie viewing. Moreover, the pattern of VT−VC connectivity differences suggested coherent systems that cut across the original subdomains. We therefore used an empirically driven reorganization–guided by NeuroMark subdomains, each ICN’s condition sensitivity, and functional annotations from Neurosynth^86^–to define eight major systems (**Fig. 4C**, **figs. S5-6**). This represents an interactive scaffold for the current study rather than a proposed new taxonomy of brain networks.

Triple-network ICNs (TN-CE, Triple Network-Central Executive; TN-DM, Default Mode; TN-SA, Salience subdomain) showed especially broad VT-related changes with other networks, motivating us to group them into three “VT-related” systems based on their FNC difference patterns. Neurosynth psychological topic mapping suggested that these systems are differentially associated with language and controlled cognition, episodic memory, and default-mode processes, and affective-reward-salience functions, respectively (**fig. S6**). In parallel, comparison with canonical parcellations^87^ showed that their spatial distributions align with known large-scale networks: one system mapped primarily onto lateral frontoparietal control regions centered in inferior frontal gyrus and dorsomedial prefrontal cortex (PFC); a second integrated frontoroparietal control territories with default-mode hubs in frontal pole/medial PFC (mPFC), temporoparietal junction, posterior cingulate cortex, and precuneus; and a third was anchored in anterior insula with extensive mPFC coverage, bridging anterior default-mode subsystems with salience/“action-mode” circuitry (**Fig. 4D**). Accordingly, we refer to these three VT-related systems as a Language-Control (LC) system, a Memory-Default (MD) system, and an Affective-Action (AA) system. The remaining five systems are: Visual (VI), Paralimbic (PL), Basal Ganglia (BG), Sensorimotor (SM), and Insular-Temporoparietal (ITP) systems (full lists in **fig. S5**).

While LC and MD showed greater within-system connectivity in VC, AA showed greater within-system coherence in VT. VT also reduced coupling among the three VT-related systems themselves (LC-MD, LC-AA, MD-AA; **Fig. 4D**), suggesting weakened coupling among these control-related ensembles under VT. The largest VT > VC effects (**Fig. 4C**; **table S10**) were observed for MD-VI, AA-BG, and within-AA (Cohen’s *d* > 0.612). The largest VT < VC effects were observed for VI-PL, ITP-PL, LC-MD, and LC-AA (Cohen’s *d* < −0.718). In addition, AA showed increased connectivity with PL, BG, SM, and ITP. Together, these patterns indicate that under VT, the affective-action system connects more with affective valuation (PL), motivational control (BG), interoceptive/mentalizing (ITP), and sensorimotor (SM) systems, while its coupling with executive and default-mode control (LC, MD) is reduced. This reorganization is consistent with a shift from an integrated, narrative-focused configuration toward a state that emphasizes threat monitoring and action readiness.

### Clustering of FNC contrasts identifies CXCL8-responsive network pattern

To identify sets of network connections that shared similar VT−VC changes and test their association with cytokine responses, we applied *k*-means clustering to the FNC contrast for the 659 Bonferroni-significant ICN pairs, resulting in 12 connectivity clusters based on optimized silhouette scores (**table S11**; *n* = 88). For each participant and cluster, we computed a centroid score that represents the mean-centered VT−VC connectivity contrast (ΔFNC) averaged across all pairs in that cluster, indexing how strongly an individual expressed each connectivity pattern. We then related these neural scores to cytokine responses, using the VT−VC fold change contrast (ΔFC) for each cytokine (positive ΔFC = larger post-scan increase or smaller decrease, after VT than VC). In linear models controlling for sex, age, and randomized condition order, only cluster 7 significantly predicted the CXCL8 ΔFC (*β* = 14.43, *P* = 0.004, *n* = 55; Bonferroni-corrected across 12 clusters; **Fig. 5**). No other clusters significantly predicted ΔFC for the cytokines (**table S12**).

**Figure 5.**
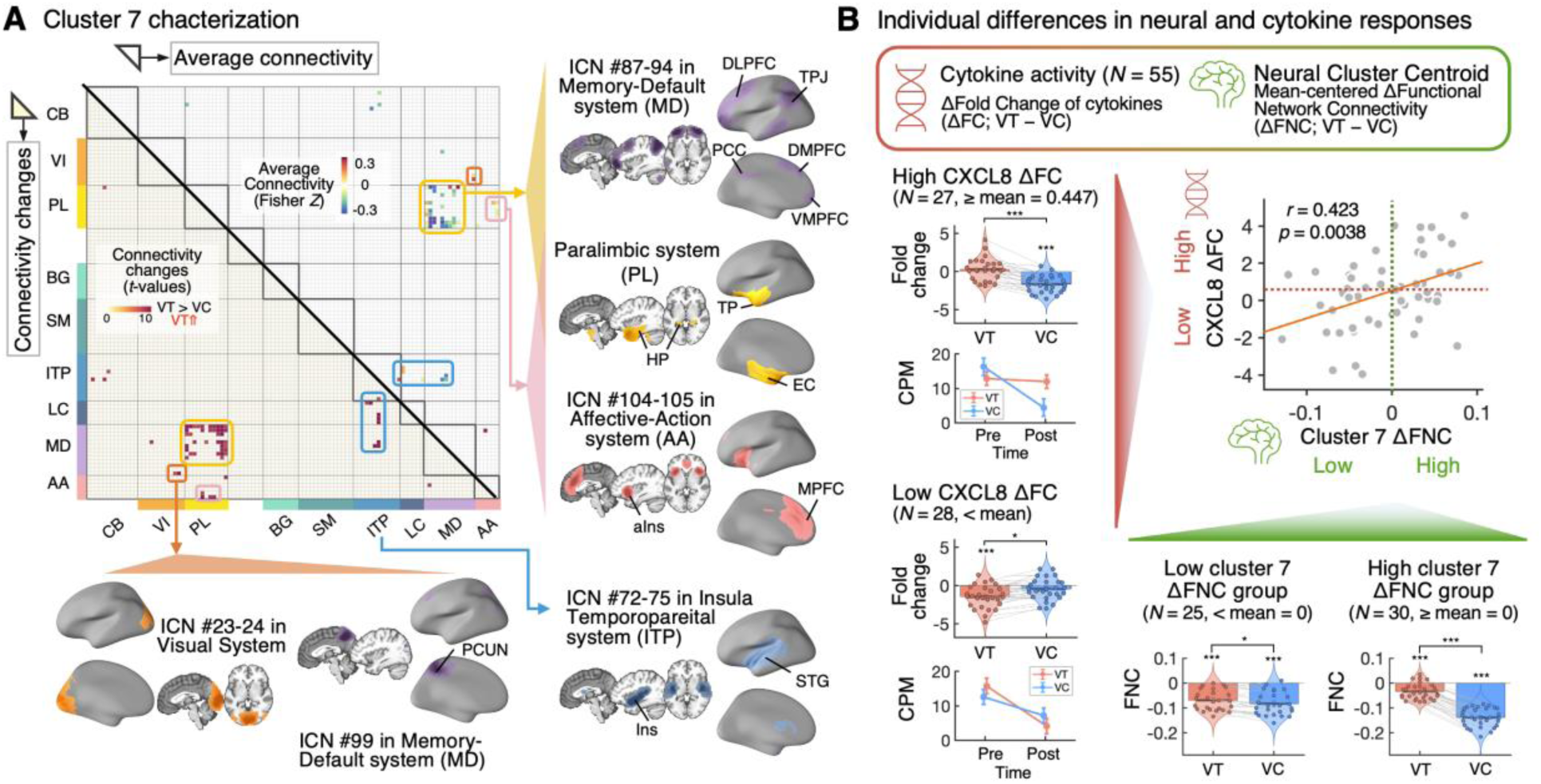
FNC cluster 7 tracks CXCL8 reactivity. (**A**) K-means clustering of VT−VC FNC contrasts across the 659 Bonferroni-significant ICN pairs identified 12 clusters, and cluster 7, which was associated with CXCL8 responses, is shown here. Lower triangle: edge-wise *t*-values (VT−VC), indicating VT > VC effects. Upper triangle: mean FNC across these edges, revealing baseline anti-correlation between PL and MD. Cluster 7 is characterized by stronger coupling between visual cortex (ICNs #23-24 in VI) and precuneus (PCUN; ICN #99 in MD), increased ITP (ICNs #72-75 in ITP) coupling with LC/MD, and weakened negative coupling between PL-MD and MD-ITP. Together, these edges index reduced coupling between paralimbic, self-referential, and control systems, and tighter embedding of visual input into default-mode hubs during VT. Abbreviation: DLPFC, dorsolateral prefrontal cortex; TPJ, temporoparietal junction; PCC, posterior cingulate cortex; DMPFC, dorsomedial prefrontal cortex; VMPFC, ventromedial prefrontal cortex; TP, temporal pole; EC, entorhinal cortex; HP, hippocampus; MPFC, medial prefrontal cortex; aIns, anterior insular cortex; Ins, Insular cortex; STG, superior temporal gyrus; PCUN, precuneus. (**B**) Cluster 7 expression predicts CXCL8 reactivity. Scatter plot shows association between participant-level cluster 7 centroid scores (ΔFNC, VT−VC; x-axis, mean-centered) and CXCL8 Δfold change (ΔFC, VT−VC; y-axis; *β* = 14.43, *P* = 0.004, *n* = 55; Bonferroni-corrected across 12 clusters), with fitted regression (*r* = 0.423, *P* = 0.004). Higher cluster 7 expression corresponds to larger VT-related increases of CXCL8. Violin and line plots show condition-wise distributions for participants stratified by ΔFC (high vs low; red triangle) and by ΔFNC (positive vs negative centroid; green triangle). Thus, individuals whose networks show greater reduction in PL-MD anticorrelation and stronger VI-MD/ITP-LC coupling in VT also show stronger VT-related increases of CXCL8 in VT.

Cluster 7 captured a coordinated pattern of VT-related connectivity increases (**Fig. 5A**). Edges in this cluster predominantly showed increased connectivity in VT between: (1) PL and MD; (2) PL and anterior insula/mPFC (in AA); (3) insular/superior temporal regions (in ITP) and LC/MD; and (4) VI and the precuneus (in MD). Thus, higher cluster-7 scores reflect reduced anti-correlation between paralimbic and default/action systems (PL-MD/AA), and stronger connectivity between interoceptive/mentalizing ITP nodes and control network (ITP-LC), as well as tighter coupling between visual input and a core self-referential hub (VI-MD).

Participants who more strongly expressed this connectivity pattern showed larger VT−VC differences in CXCL8 expression. Higher cluster 7 ΔFNC scores were associated with greater CXCL8 ΔFC. To further characterize this relationship, we stratified participants by CXCL8 ΔFC (high ≥ mean 0.447 vs. low < mean) and by cluster 7 centroid (positive vs. negative ΔFNC; **Fig. 5B**). In the high ΔFC group (*n* = 27), CXCL8 decreased significantly after VC but not after VT, replicating the full-sample pattern (**Fig. 3C**). In contrast, in the low ΔFC group (*n* = 28), CXCL8 decreased after VT but not after VC. These opposite trends across groups indicate that higher ΔFC reflects a relative increase of CXCL8 under VT with little change in VC, whereas lower ΔFC reflects a decrease in VT with stable levels in VC. Although the high-ΔFC group mirrored the overall group-level trajectory, the divergence between high- and low-ΔFC subgroups underscores substantial individual differences in CXCL8 responses to vicarious trauma.

For the neural measure, both centroid groups showed negative mean connectivity among cluster 7 edges in both conditions, but participants with a positive centroid (ΔFNC; *n* = 30) showed significantly weaker anticorrelation in VT than in VC. Thus, individuals who showed a VT-related increase of CXCL8 also showed a VT-related reduction in anti-correlation between paralimbic and Memory-Default systems and enhanced coupling between visual and Memory-Default networks and between insular/temporal ITP and Language-Control systems. This identifies a specific large-scale connectivity configuration whose VT-evoked change tracks CXCL8 chemokine responses.

## Discussion

Vicarious trauma can evoke moral injury and long-term health risks, yet its physiological mechanisms in humans remain poorly characterized under ecologically realistic conditions. Although moral injury and vicarious trauma are associated with PTSD, depression, suicidality, and chronic disease^6,7,40,41^, prior work has not provided causal, mechanistic evidence that morally salient vicarious trauma in humans reorganizes neural and immune consequences. Guilt- and shame-evoking stressors are known to heighten inflammation^43,44^, but it has been unclear how real-world vicarious trauma engages the neural system and programs innate immune cells such as monocytes. Here, we addressed this gap using a within-subject documentary paradigm in which participants viewed a 30-minute film depicting industrial animal slaughter (VT) and, on a separate day, a matched film of positive human-animal interactions (VC) while we recorded fMRI, autonomic signals, and monocyte transcriptomes. Relative to VC, VT induced persistent sympathetic arousal, an anticipatory shift in monocytes toward a pro-inflammatory transcriptional profile (upregulation of IL1B and CXCL8, downregulation of TGFB1), and a large-scale reconfiguration of multiple neural networks. Specifically, VT reduced integration within and between Language-Control (LC) and Memory-Default (MD) systems and increased coupling of the Affective-Action system (AC) with insular, striatal, and sensorimotor regions. Moreover, a VT > VC connectivity pattern linking paralimbic and MD systems tracked higher CXCL8 after VT, tying neural network shifts to an innate immune chemokine associated with neutrophil mobilization. Together, these convergent autonomic, immune, and neural changes under naturalistic viewing suggest a centrally coordinated, anticipatory defensive state during morally salient vicarious trauma.

Beyond general distress, vicarious trauma in this paradigm elicited a moral/shame-related emotion dimension that tracked sustained elevations in heart rate. Participants reported these moral/social emotions, captured by EmoPC2, distinguishing this study from prior vicarious distress studies that rely on decontextualized or fictional stimuli. Autonomically, VT (vs. VC) was associated with elevated heart rate that persisted for up to 40 minutes after movie viewing—substantially longer than is typically observed in typical laboratory stress effects^25,88^. Moreover, movie-induced negative valence (EmoPC1) predicted elevated heart rate, especially for clips that elicited stronger empathic engagement (EmoPC2), indicating that moral emotional intensity amplified sympathetic activation. These profiles align with animal and human work showing that observing others’ distress elevates heart rate^12^, corticosterone^9,11,12,34^, and skin conductance^29,35^ and underscore the potential physiological costs of vicarious trauma and motivating tests of downstream immune programming.

Extensive work has linked stress to immune changes, but most human studies rely on circulating hormones or cytokines, which cannot reveal which cell types are responsible or how^26,65–69^. Because these cytokines are produced by multiple immune populations and stress redistributes cells across blood and tissues, protein-level assays blur changes in cell numbers with changes in cell-intrinsic programming^70–72^. To isolate cell-specific threat responses, we focused on circulating monocytes whose transcriptional profile provides an early readout of inflammatory priming and neutrophil-recruiting capacity^74^. RNA-seq provided sensitive characterization of monocyte programming, capturing anticipatory shifts that can precede detectable changes in circulating proteins and providing whole-transcriptome coverage beyond the targets reported here^79–81^. Within this framework, we selected four a priori cytokine/chemokine genes for focused analysis based on (1) robust monocyte expression, reliability in our dataset, and established relevance to stress-linked inflammation: IL1B, CXCL8, TNF, and TGFB1 (details in Materials and Methods).

IL-1β is a key “threat amplification” cytokine that primes and propagates inflammatory responses in the brain and periphery and is upregulated by acute and chronic psychosocial stress^26,27,69,70,84,85,89,90^. IL-8 drives neutrophil recruitment and activation^75,91^ and is often elevated following stress exposure, consistent with mobilization of innate defenses^84,85^. TNF-α is a canonical pro-inflammatory mediator^92^ whose increase is a common feature of chronic stress and stress-related disorders^26,27,84,85,90^. In contrast, TGF-β1 serves as a central counter-regulatory cytokine that constrains excessive inflammation and promotes resolution^93–95^. These markers are prominently produced by circulating monocytes^80,85^ and are modulated by stress-responsive pathways, including NF-κB and SMAD/TGF-β signaling^95–97^. Therefore, this panel indexes both pro-inflammatory drive and regulatory control, capturing a balance that is repeatedly implicated in stress-related inflammatory phenotypes.

In this context, our results suggest that VT induced a neutrophil-recruiting, pro-inflammatory bias with weakened regulatory braking. IL1B and CXCL8—two of the three pro-inflammatory markers—increased after VT, particularly in participants with higher baseline levels, whereas TGFB1 decreased on average, indicating diminished anti-inflammatory regulatory tone under morally salient vicarious stress. These findings point to stress-amplified innate immune priming and differential immune susceptibility rather than a uniform surge. The selective IL1B/CXCL8 shifts alongside stable TNF are also informative. Classical microbial triggers such as LPS, acting via TLR4–MyD88 signaling, induce a broad cytokine profile that includes TNF-α, IL-1β, and CXCL8, whereas inflammasome-centric signaling more selectively drives IL-1β with relatively weaker TNF-α induction. The present pattern—IL1B/CXCL8 changes without TNF—therefore points to a nuanced, sterile, neurogenic inflammatory state in monocytes rather than a generalized cytokine surge.

The observed cytokine shifts are unlikely to reflect simple peripheral reactions to stress, but rather a brain-engaged anticipatory program, supported by extensive animal and human studies. The brain predictively tunes immunity via autonomic innervation of lymphoid organs^98–100^, stress-evoked release of noradrenaline and glucocorticoids, which facilitate monocyte production and mobilize from bone marrow and spleen^101–103^, and vagal cholinergic anti-inflammatory reflex that actively constrains inflammation, demonstrating bidirectional neural control^104,105^. Beyond these hardwired pathways, higher-order brain functions can program immune responses: classic conditioning studies showed that learned cues can change immunity^106,107^, and circuit-level work has identified causal ensembles^108^, such as insular cortex populations whose reactivation triggers inflammation without any pathogen^109^. Reward circuitry activation can enhance systemic immunity via sympathetic outputs^110^, and specific forebrain circuits can drive inflammation^111,112^. In humans, convergent work on neurovisceral integration and emotional awareness^113,114^, default-mode and control network decoupling in stress-related psychopathology^115^, and brain networks coupling between heart rate variability and emotion-regulation circuitry^116,117^ underscores that prefrontal, cingulate, insular, and default-mode systems jointly organize interoception, autonomic regulation, and self-related processing. Together, these findings establish a mechanistic basis for the causal brain-to-immune coding of anticipated threats, situating our transcriptomic findings within a plausibly brain-driven program and motivating our neural network analyses.

Building on these neuroimmune findings, we asked whether VT reorganizes large-scale brain networks and whether those changes relate to immune priming. To do so, we combined extended naturalistic fMRI with an individualized network-mapping approach that fit reliable network templates to each participant’s 30-minute VT and VC runs. This strategy leverages long, continuous viewing to improve within-person reliability^118,119^ while taking account of individual differences in functional anatomy, providing participant-specific network configurations that we could directly relate to monocyte transcriptional responses.

Our findings reveal that vicarious trauma provokes a significant shift in large-scale brain dynamics, transitioning the system from a mode of integrated experience to one of anticipatory defense. Within the triple-network^120,121^, which allocates resources between internal and external demands, among the Affective-Action (AA, corresponding to the Salience network), Memory-Default (MD, Default-Mode network), and Language-Control (LC, Frontoparietal network), VT increased segregation (greater anti-correlation) between salience/“action-mode” (AA) and frontoparietal control (LC) systems, indicating a state in which threat detection is prioritized over flexible executive control—a trade-off that is adaptive for survival but costly for ongoing cognitive goals^122–125^. This state was not merely attentional but also deeply embodied. The AA network became tightly coupled with a cortico-subcortical action circuit–including the hippocampus, basal ganglia, insula, and sensorimotor regions–suggesting translation of interoceptive threat signals into action readiness. Conversely, the VC fostered a more integrated state, with stronger coupling between AA and MD systems, a pattern consistent with narrative absorption and social cognition, in which the salience of social-emotional cues guides self-referential processing^126^. Together, these results reveal that vicarious trauma reorganizes whole-brain networks to implement an anticipatory threat response, dynamically balancing vigilant reactivity against immersive integration.

Critically, this neural shift into an anticipatory defense mode was directly linked to innate immune priming. A specific, multivariate connectivity pattern—reflecting reduced decoupling between paralimbic and Memory-Default systems and stronger integration of visual, paralimbic, insular/temporal areas—predicted individual differences in CXCL8 upregulation (VT > VC Δfold change). This pattern suggests that visceral-affective states exerted greater influence on self-related processing, consistent with a view where emotional experiences are generated from the integration of interoceptive signals with higher-order appraisal and conceptualization^127,128^. Concurrently, the witnessed suffering was processed as part of one’s own embodied experience, evidenced by stronger connectivity between the visual cortex and the precuneus, a key region for self-awareness and embodied perspective-taking^129–131^. Collectively, this network reconfiguration integrated visceral, perceptual, and affective information into the core of the self-regulatory neural systems, creating a state primed for an embodied threat response^132^. These findings provide a direct link between the brain’s adoption of an anticipatory, threat-oriented state and the mobilization of a specific innate immune pathway designed to counter the physical harm it predicts.

This study has limitations that guide future work. Although our naturalistic films produced robust effects across systems, they focused on nonhuman suffering; analogous paradigms using human trauma are needed to test generalizability. The VC condition served as an affiliative, content-matched control rather than a true neutral baseline, so future designs should consider perceptually matched neutral content to dissociate valence, arousal, and moral salience. Our findings suggest that individuals routinely exposed to vicarious trauma–such as those working in food production, healthcare, emergency responses, or military roles–may be at risk for cumulative alterations in brain and immune function. Longitudinal and occupational cohort studies will be crucial to determine whether the anticipatory patterns observed here accumulate to influence mental and physical health over time. Future work should also examine moderators such as moral conviction about animals, dietary practices, prior trauma, sleep, and medications, all of which can shape sympathetic tone and immune programming and may help identify individuals most vulnerable to sustained allostatic responses. Denser temporal sampling, combined with proteomic and cellular phenotyping, would further clarify the temporal cascade by which vicarious trauma exerts its psychobiological effects.

In sum, this study shows that vicarious trauma is sufficient to shift the brain into an anticipatory defensive mode and to prime innate immune cells towards a neutrophil-recruiting, pro-inflammatory state. Large-scale network reconfiguration integrating visual, self-referential, and paralimbic-visceromotor systems predicted CXCL8 upregulation in monocytes during VT, directly linking embodied moral experience to a specific inflammatory pathway. These findings demonstrate that vicarious trauma can reorganize human brain-body systems in ways that plausibly accumulate into long-term health risk and provide a mechanistic framework for understanding the biological embedding of moral injury.

## Acknowledgments

We thank Elizabeth C. Tremmel, Bob J. Yang, and the Cognitive and Affective Neuroscience Lab for insightful comments and discussions; Terry J. Sackett, and Courtney Rogers for technical support in data collection; Yaroslav O. Halchenko for MRI data curation. This manuscript was proofread and checked for grammar using ChatGPT and Gemini. All content was reviewed and verified by the authors. The work was supported by NIMH R01 MH116026 (T.D.W), MH-R37-076136 (T.D.W), R01EB026549 (T.D.W), Animals Australia for a Kinder World (T.D.W), and NIH R35 HL155458 (C.V.J.). Bulk RNA sequencing was carried out in the Genomics and Molecular Biology Shared Resource (RRID:SCR_021293) at Dartmouth, which is supported by NCI Cancer Center Support Grant 5P30CA023108 and NIH S10 (1S10OD030242) awards.

## Author contributions

Conceptualization: BKL, MCK, CVJ, TDW

Data curation: BKL, MCK, DW

Formal analysis: BKL, XL, TDW

Funding acquisition: CVJ, TDW

Investigation: BKL, MCK, DW, FWK, XL

Project administration: BKL, TDW

Resources: BKL, MCK, FWK, XL, CVJ

Supervision: TDW Visualization: BKL, CVJ

Writing – original draft: BKL, TDW

Writing – review & edition: BKL, MCK, XL, CVJ, TDW

## Data and materials availability

All preprocessed data and code used in this study are available on Zenodo at [https://zenodo.org/records/17834309]. All other data are available in the main text or the supplementary materials.

## Supplementary Materials

Methods Figs. S1 to S7

Tables S1 to S12

