## Supplementary Materials for "Vicarious trauma primes innate immunity and reconfigures human brain networks"

**The PDF file includes:**

Materials and Methods
Figs. S1 to S7
Tables S1 to S12

### Methods

#### *Study*

This study was conducted from February to November 2022 at the Dartmouth Brain Imaging Center (DBIC) at Dartmouth College, NH, USA. The functional Magnetic Resonance Imaging (fMRI) experiment was conducted alongside physiological activity recording, self-report measurements, behavioral assessments, and blood samples for immune assays. Participants visited the lab on two separate days for the fMRI sessions (mean interval = 9.88 ± 9.48 days). Sessions were conducted in the morning (before noon) to minimize variability related to diet and the circadian rhythms of cytokines. The institutional review board of Dartmouth College approved the study, and all participants provided written consent and received compensation for their participation. Stimulus presentation and behavioral data acquisition were performed using Psychtoolbox-3 (http://psychtoolbox.org/), MATLAB (MathWorks, Natick, MA), and PsychoPy (https://psychopy.org/[133]).

#### *Participants*

Eighty-eight participants (age = 24.38 ± 7.97 years, 48 females) from the Upper Valley community in and around Hanover, NH, participated in the fMRI study. Before they visited the DBIC, they completed a pre-visit self-report survey via Qualtrics. Preclusion criteria included self-reported psychiatric, neurological, or systemic disorders as well as MR contraindications. Participants also self-reported race and ethnicity to characterize the sample. To minimize potential confounds on blood gene expression and control for circadian variation, all experimental sessions were conducted in the morning before participants ate lunch.

#### *Self-report measurements*

Participants responded to a set of self-report questionnaires during the pre-visit online survey. The personality measures include Interpersonal Reactivity Index (IRI) which assesses empathic traits[134].

#### *fMRI experiments*

##### *Movie-viewing runs*

Two 30-minute animal documentaries, professionally produced and assembled from real archival footage, were presented during movie viewing runs. The Vicarious Trauma (VT) documentary showed industrial handling, transport, and slaughter of farmed animals common in Western food production, as well as animal testing procedures, emphasizing scenes of suffering and harm. The Vicarious Community (VC) documentary showed positive human-animal interactions, including animal rescue and affectionate interaction with farm animals, highlighting animal intelligence, social behavior, and affiliative human contact. Both documentaries include spoken narration but no background music.

Each participant viewed both documentaries on separate days in a counterbalanced order. In each session, participants watched one documentary presented across three fMRI runs. Each documentary was divided into nine clips, with each run containing three 12-minute clips. After each clip, participants rated their emotional responses on seven dimensions: “warm and tender,” “joyful,” “inspired or uplifted,” “ashamed,” “sad,” “horrified,” and “disgusted”. We included both positive and negative emotions, spanning higher-level and lower-level emotional categories[135]. The order of the emotion ratings was randomized on each trial. Ratings were collected using a visual rating bar, and participants reported their response by selecting a point on the bar with an MR-compatible trackball using their right hand. In total, participants provided nine trials of emotion ratings per documentary, each covering all seven emotion categories.

*Resting-state runs*

After the movie-viewing runs, participants completed a ten-minute resting-state session, during which they were instructed to relax.

***Blood collection***

Peripheral blood was collected via venipuncture by certified phlebotomists before and after the fMRI scan in a behavioral experiment room located in the same building as the MRI scanner ($n$ = 70). A total of 8 mL of peripheral blood was collected using BD Vacutainer Mononuclear Cell Preparation Tubes with Sodium Heparin. The samples were stored at 4°C immediately after collection.

***Blood sample inclusion***

Peripheral blood was collected from 70 participants. Each participant was expected to contribute four monocyte samples (pre- and post-scan across two sessions). To ensure complete data for within-subject modeling, participants missing any of the four samples were excluded. Exclusions reflected issues arising during blood collection, sample processing, or RNA-seq quality control. After these criteria were applied, 55 participants had complete, high-quality monocyte RNA-seq data and were retained for all analyses.

***Sample processing and cell isolation***

All samples were processed by the same person (BKL) in the Jakubzick lab at Dartmouth College. At room temperature, the whole blood samples were centrifuged at 2000 RCF (Relative Centrifugal Force) for 20 minutes. Following centrifugation, 1.4 mL of plasma from the uppermost layer was collected using a Pasteur Pipette, mixed with 0.1 mL cOmplete protease inhibitor (Roche) solution, and stored at -80°C until further processing. The remaining mononuclear cells and platelets were washed with MACS buffer, consisting of 1X Ca2+/Mg2+ free Phosphate Buffered Saline (PBS, Corning), 0.5% Bovine Serum Albumin (BSA, Sigma), and 1mM Ethlenediaminetetraacetic acid (EDTA, Invitrogen). Approximately 60% of the cells were used for monocyte enrichment, and the remaining ~40% were analyzed via flow cytometry using specific antibody panels.

Monocytes were enriched via negative selection using the EasySep Human Monocyte Isolation Kit (Stemcell). A portion of the isolated monocytes was assessed via flow cytometry to verify enrichment quality using antibody panel B. RNA was extracted from the monocytes using the RNeasy Mini Kit (Qiagen) and stored at -80°C until submission to the Genomic Core for RNA sequencing.

#### *Flow cytometry staining and analysis*

Whole blood cells and enriched monocytes were analyzed by flow cytometry. Isolated cells were stained with the following anti-human antibody panels to examine samples before and after enrichment. The panel consisted of CD88 (S5/1), CD141 (REA674), HLA-DR (L243), CD11B (ICRF44), CD16 (CB16), CD1C (AD5-8E7), and CD14 (M5E2). Panel C included CD88 (S5/1), CD20 (2H7), CD4 (SK3), CD15 (80H5), CD26 (BA5b), CD56 (B159), CD8 (SK1), and CD69 (FN50).

After 1 hour of antibody staining, DAPI was added to exclude dead cells, and samples were analyzed by flow cytometry. Data were acquired using a BD Symphony and analyzed with FlowJo software. Monocyte enrichment efficiency averaged 93.04% ± 9.88% (mean ± SD). Sixteen of the 220 samples (from 55 participants) showing hemolysis, failed staining, or unsuccessful image acquisition were excluded from the cytometry efficiency calculation. However, because these samples met RNA-seq quality control criteria, they were retained for all downstream analyses.

#### *Whole transcriptome sequencing (RNA-seq)*

##### *Library construction*

RNA sequencing was performed by the Genomics and Molecular Biology Shared Resource (GMBSR) at the Geisel School of Medicine at Dartmouth. RNAs were quantified by qubit (Thermo Fisher), and quality was assessed using a Fragment Analyzer (Agilent). Using 100 ng of RNA as input, samples were first ribo-depleted to remove rRNA using FastSelect HMR probes (Qiagen) and then prepared using the Kapa RNA HyperPrep kit (Roche) following the manufacturer’s instructions, with 12 PCR cycles for final library amplification. Libraries were quantified by Qubit and Fragment Analyzer and pooled for sequencing on a NextSeq2000 (Illumina) using paired-end 50bp reads, targeting 30 million reads per sample.

##### *Read alignment*

Sequencing reads were aligned to hg38 (UCSC human genome 38) using STAR, and gene-level counts were quantified with featureCounts (Subread package). Quality control was performed using FastQC to ensure sufficient sequencing depth, with total read counts exceeding 15 million per sample. All retained samples had more than 21 million mapped reads.

##### *Expression quantification and normalization*

Raw read counts were normalized in two ways to meet different analytic goals. First, we calculated transcripts per million (TPM), which adjusts for both gene length and library size and therefore supports comparisons across genes within a sample (and across samples for broad abundance checks). We used TPM to screen a priori cytokine targets with robust expression and to validate monocyte enrichment with marker panels that require cross-gene marker comparability (**Fig. 3B**). Second, we calculated counts per million (CPM), which adjusts only for library size and preserves the appropriate variance structure for within-gene comparison.

Because gene length is constant in our within-subject repeated-measures models, CPM was used for all downstream analysis of condition effects. Thus, TPM was used for target selection and cell-type validation, and CPM for all inferential analyses.

Counts per million (CPM) values were computed using the EdgeR package[136]. Raw gene counts were first normalized for library size using the Trimmed Mean of M-values (TMM) method implemented in the calcNormFactors() function. Normalized counts were then converted to CPM values using the cpm() function, which provides a stabilized, library-size–adjusted expression measure suitable for within-gene comparisons across conditions and sessions. For fold-change estimates for each condition, $\log_2$ fold change was calculated from pre- and post-scan CPM values as follows: $\log_2 \frac{Post+0.146}{Pre+0.146}$, with a 0.146 pseudocount added to accommodate zero values.

#### *Targeted assessment of cytokines*

We selected cytokines a priori based on two criteria: established relevance to psychosocial stress biology and robust transcription in circulating monocytes in our bulk RNA-seq data (TPM > 10). Guided by key reviews and meta-analyses of stress-responsive immune markers[26,27,69,84,85,90], we initially considered eleven cytokines: IL-1β, IL-6, CRP, TNF-α, CXCL8, IFN-γ, IL-12, IL-2, TGF- β1, IL-4, and IL-10. On theoretical grounds of cellular origin, several of these markers are not primarily produced by monocytes: IL-2, IL-4, IFN-γ, and IL-12 are predominantly lymphocyte-derived; CRP is hepatocyte-derived; and IL-6 and TGF-β have multiple sources of cells. Consistent with this expectation, IL2, IL4, IFNG, IL12, CRP, and IL6 showed low monocyte expression in our dataset (mean TPM ranges 0.06 to 3.20), below our inclusion threshold (TPM > 10).

| Gene | IL1B | IL2 | IL4 | IL6 | CXCL8 | IL10 |
|---|---|---|---|---|---|---|
| mean | 34.81 | 0.06 | 3.20 | 0.31 | 15.70 | 1.61 |
| STD | 15.40 | 0.13 | 2.55 | 0.52 | 17.18 | 0.84 |
| Gene | IL12A | IL12B | IFNG | CRP | TGFB1 | TNF |
| mean | 1.89 | 0.06 | 1.87 | 0.07 | 510.83 | 22.51 |
| STD | 1.23 | 0.21 | 2.02 | 0.16 | 227.39 | 6.67 |

**Table. TPM levels of candidate cytokine targets**

We therefore focused on IL1B (ENSG00000125538), CXCL8, TNF(ENSG00000232810), and TGFB1 (ENSG00000105329) ($n$ = 55). Among the TGF-β isoforms, we selected TGF-β1 because it is the dominant isoform expressed by monocytes and the anti-inflammatory cytokines inhibited by stress hormones[85,93,95]. Targets were defined prior to all downstream analyses, and CPM values were used for all subsequent modeling.

#### *Mixed-effects analysis of cytokine expression*

To examine the effects of condition and its interaction with baseline gene expression on post-scan gene expression levels, we fitted linear mixed-effects models using MATLAB 'fitlme' function. The outcome variable was post-scan gene expression, and the predictors included mean-centered baseline expression (Pre), experimental condition, their interaction, randomized condition order, sex, age, and batch. The models included participant-specific random intercepts and random slopes for condition to account for within-subject correlation and individual variability in condition effect, with partial pooling improving stability and precision of fixed-effect estimates[137–139]. The model formula was: Post ~ Condition * Pre + Condition Order + Sex + Age + Batch + (1 + Condition |Subject). Models were estimated using restricted maximum likelihood (REML), and degrees of freedom for fixed effects were calculated using the Satterthwaite approximation via the 'fixedEffects' function.

#### *Physiological activity recording and preprocessing*

During the fMRI scan, heart rate and skin conductance were recorded using photoplethysmography (PPG) and Electrodermal activity (EDA), respectively. PPG was recorded by attaching an MR-compatible PPG transducer (Biopac Systems, Goleta, CA) to the left thumb. EDA was recorded using MR-compatible electrodes (Biopac Systems, Goleta, CA) placed on the left index and middle fingers. Both signals were sampled at 2,000Hz.

PPG preprocessing included band-pass filtering (0.5-5Hz) and a comb filter at the repetition frequency of 1/1.3 Hz (TR) to remove MR-induced noise. Peak detection and inter-beat interval (IBI) calculations were performed using the PhysIO Toolbox (https://www.nitrc.org/projects/physio/). IBI data were downsampled to 1Hz. Implausible data points were identified using two criteria: (1) IBIs shorter than 0.5 seconds (equivalent to 120 beats per minute), given that participants were lying in the scanner, and (2) IBI differing from the immediately preceding IBI by more than three times the participant-specific standard deviation of successive differences. Outliers were replaced via linear interpolation between adjacent values. Heart rate (beats per minute, BPM) was computed by dividing 60 seconds by the IBI.

EDA data were downsampled to 10Hz first. For outlier detection, we computed the participant-specific standard deviation of successive EDA differences, as PPG. Any sample differing from the previous sample by more than three times this value was labeled an outlier and replaced via linear interpolation. The signal was then low-pass filtered at 1Hz and is hereafter referred to as the skin conductance response (SCR) signal.

After the fMRI scan, a five-minute resting physiology recording was conducted in a behavioral experiment room to measure heart rate and skin conductance. To avoid the potential effects of venipuncture on physiology, recording took place while participants were seated before the second blood draw. Heart rate was recorded with both PPG and electrocardiography (ECG). Three ECG electrodes were placed on the right and left clavicles and the middle abdomen. The PPG transducer and EDA electrodes were positioned identically to the fMRI scan setup on the left hand. Preprocessing was identical to the in-scanner pipeline, except that the comb filtering for IBI used to remove MRI noise was omitted. ECG data preprocessing followed the same steps as the PPG data.

Participants with incomplete recordings or poor signal quality were excluded. Only those with complete data, including in-scanner recording and post-scan resting recording, were retained for analysis. Consequently, heart rate data from 3 participants and skin conductance data from 16 participants were excluded.

#### *MRI acquisition*

FMRI data were acquired on a 3T Siemens MAGNETOM Prisma MRI scanner with 32-channel parallel imaging at the Dartmouth Brain Imaging Center. We acquired structural (T1 imaging) and functional images (multi-echo multiband planar imaging).

Structural images were acquired using a high-resolution T1-weighted magnetization-prepared rapid acquisition gradient echo (MPRAGE) imaging sequence (TR = 2000 ms, TE = 2.11 ms, Flip angle = 8°, FOV = 256 × 256 mm, number of slices = 224, voxel size = 0.8 mm isotropic, Multiband Acceleration Factor = 3, Slice orientation = Sagittal, iPAT mode = GRAPPA, Echo spacing = 7.2ms, Bandwidth = 240 Hz/px).

Functional images were acquired using a multi-echo multiband T2*-weighted sequence (TR = 1300 ms, TE = 13.20, 31.45, 49.7 ms, Flip angle = 60°, FOV = 240 × 240 mm, number of slices = 51, voxel size = 2.7 mm isotropic, Multiband Acceleration Factor = 3, iPAT mode = GRAPPA, Slice orientation = Transversal, Interleaved slice acquisition, Phase encoding direction = anterior to posterior, Echo spacing = 0.49ms, Bandwidth = 2778 Hz/px). We discarded the first five images, along with those automatically removed by the MRI scanner, to ensure the signal reached a steady state[140].

#### *MRI data curation*

MRI protocols were renamed to follow the ReproIn naming convention, enabling robust and automated conversion into BIDS format using HeuDiConv v.0.9.0[141]. For sequences not initially named according to ReproIn, a remapping was provided to facilitate conversion. Subject IDs were anonymized and assigned sequentially within the study, ranging from 1 to 88, with zero-padding to ensure two-digit formatting (e.g., sub-01).

#### *fMRIPrep*

FMRI data were preprocessed using fMRIPrep 23.1.3[142]. The following description is generated from the standard fMRIPrep preprocessing boilerplate text.

Results included in this manuscript come from preprocessing performed using *fMRIPrep* 23.1.3 (https://fmriprep.org/en/23.1.3/index.html) (Esteban et al. (2019); Esteban et al. (2018); RRID:SCR_016216), which is based on *Nipype* 1.8.6 (K. Gorgolewski et al. (2011); K. J. Gorgolewski et al. (2018); RRID:SCR_002502).

*Preprocessing of B0 inhomogeneity mappings*

A total of 2 fieldmaps were found available within the input BIDS structure for this particular subject. A *B0*-nonuniformity map (or *fieldmap*) was estimated based on two (or more) echo-planar imaging (EPI) references with topup (Andersson, Skare, and Ashburner (2003); FSL None).

*Anatomical data preprocessing*

A total of 2 T1-weighted (T1w) images were found within the input BIDS dataset. All of them were corrected for intensity non-uniformity (INU) with N4BiasFieldCorrection (Tustison et al. 2010), distributed with ANTs (version unknown) (Avants et al. 2008, RRID:SCR_004757). The T1w-reference was then skull-stripped with a *Nipype* implementation of the antsBrainExtraction.sh workflow (from ANTs), using OASIS30ANTs as target template. Brain tissue segmentation of cerebrospinal fluid (CSF), white-matter (WM) and gray-matter (GM) was performed on the brain-extracted T1w using fast (FSL (version unknown), RRID:SCR_002823, Zhang, Brady, and Smith 2001). An anatomical T1w-reference map was computed after registration of 2 T1w images (after INU-correction) using mri_robust_template (FreeSurfer 7.3.2, Reuter, Rosas, and Fischl 2010). Brain surfaces were reconstructed using recon-all (FreeSurfer 7.3.2, RRID:SCR_001847, Dale, Fischl, and Sereno 1999), and the brain mask estimated previously was refined with a custom variation of the method to reconcile ANTs-derived and FreeSurfer-derived segmentations of the cortical gray-matter of Mindboggle (RRID:SCR_002438, Klein et al. 2017). *Grayordinate* "dscalar" files (Glasser et al. 2013) containing 91k samples were also generated using the highest-resolution fsaverage as an intermediate standardized surface space. Volume-based spatial normalization to two standard spaces (MNI152NLin2009cAsym, MNI152NLin6Asym) was performed through nonlinear registration with antsRegistration (ANTs (version unknown)), using brain-extracted versions of both T1w reference and the T1w template. The following templates were were selected for spatial normalization and accessed with *TemplateFlow* (23.0.0, Ciric et al. 2022): *ICBM 152 Nonlinear Asymmetrical template version 2009c* [Fonov et al. (2009), RRID:SCR_008796; TemplateFlow ID: MNI152NLin2009cAsym], *FSL's MNI ICBM 152 non-linear 6th Generation Asymmetric Average Brain Stereotaxic Registration Model* [Evans et al. (2012), RRID:SCR_002823; TemplateFlow ID: MNI152NLin6Asym].

*Functional data preprocessing*

For each of the 14 BOLD runs found per subject (across all tasks and sessions), the following preprocessing was performed. First, a reference volume and its skull-stripped version were generated by aligning and averaging 2 single-band references (SBRefs). Head-motion parameters with respect to the BOLD reference (transformation matrices, and six corresponding rotation and translation parameters) are estimated before any spatiotemporal filtering using mcflirt (FSL , Jenkinson et al. 2002). The estimated *fieldmap* was then aligned with rigid-registration to the target EPI (echo-planar imaging) reference run. The field coefficients were mapped on to the reference EPI using the transform. BOLD runs were slice-time corrected to 0.602s (0.5 of slice acquisition range 0s-1.21s) using 3dTshift from AFNI (Cox and Hyde 1997, RRID:SCR_005927). A T2★ map was estimated from the preprocessed EPI echoes, by voxel-wise fitting the maximal number of echoes with reliable signal in that voxel to a monoexponential signal decay model with nonlinear regression. The T2★/S0 estimates from a log-linear regression fit were used for initial values. The calculated T2★ map was then used to optimally combine preprocessed BOLD across echoes following the method described in (Posse et al. 1999). The optimally combined time series was carried forward as the *preprocessed BOLD*. The BOLD reference was then co-registered to the T1w reference using bbregister (FreeSurfer) which implements boundary-based registration (Greve and Fischl 2009). Co-registration was configured with nine degrees of freedomto account for distortions remaining in the BOLD reference. First, a reference volume and its skull-stripped version were generated using a custom methodology of *fMRIPrep*. Several confounding time-series were calculated based on the *preprocessed BOLD*: framewise displacement (FD), DVARS and three region-wise global signals. FD was computed using two formulations following Power (absolute sum of relative motions, Power et al. (2014)) and Jenkinson (relative root mean square displacement between affines, Jenkinson et al. (2002)). FD and DVARS are calculated for each functional run, both using their implementations in *Nipype* (following the definitions by Power et al. 2014). The three global signals are extracted within the CSF, the WM, and the whole-brain masks. Additionally, a set of physiological regressors were extracted to allow for component-based noise correction (*CompCor*, Behzadi et al. 2007). Principal components are estimated after high-pass filtering the *preprocessed BOLD* time-series (using a discrete cosine filter with 128s cut-off) for the two *CompCor* variants: temporal (tCompCor) and anatomical (aCompCor). tCompCor components are then calculated from the top 2% variable voxels within the brain mask. For aCompCor, three probabilistic masks (CSF, WM and combined CSF+WM) are generated in anatomical space. The implementation differs from that of Behzadi et al. in that instead of eroding the masks by 2 pixels on BOLD space, a mask of pixels that likely contain a volume fraction of GM is subtracted from the aCompCor masks. This mask is obtained by dilating a GM mask extracted from the FreeSurfer's *aseg* segmentation, and it ensures components are not extracted from voxels containing a minimal fraction of GM. Finally, these masks are resampled into BOLD space and binarized by thresholding at 0.99 (as in the original implementation). Components are also calculated separately within the WM and CSF masks. For each CompCor decomposition, the *k* components with the largest singular values are retained, such that the retained components' time series are sufficient to explain 50 percent of variance across the nuisance mask (CSF, WM, combined, or temporal). The remaining components are dropped from consideration. The head-motion estimates calculated in the correction step were also placed within the corresponding confounds file. The confound time series derived from head motion estimates and global signals were expanded with the inclusion of temporal derivatives and quadratic terms for each (Satterthwaite et al. 2013). Frames that exceeded a threshold of 0.9 mm FD or 1.5 standardized DVARS were annotated as motion outliers. Additional nuisance timeseries are calculated by means of principal components analysis of the signal found within a thin band (*crown*) of voxels around the edge of the brain, as proposed by (Patriat, Reynolds, and Birn 2017). The BOLD time-series were resampled into standard space, generating a *preprocessed BOLD run in MNI152NLin2009cAsym space*. First, a reference volume and its skull-stripped version were generated using a custom methodology of *fMRIPrep*. The BOLD time-series were resampled onto the left/right-symmetric template "fsLR" (Glasser et al. 2013). *Grayordinates* files (Glasser et al. 2013) containing 91k samples were also generated using the highest-resolution fsaverage as intermediate standardized surface space. All resamplings can be performed with *a single interpolation step* by composing all the pertinent transformations (i.e. head-motion transform matrices, susceptibility distortion correction when available, and co-registrations to anatomical and output spaces). Gridded (volumetric) resamplings were performed using antsApplyTransforms (ANTs), configured with Lanczos interpolation to minimize the smoothing effects of other kernels (Lanczos 1964). Non-gridded (surface) resamplings were performed using mri_vol2surf (FreeSurfer).

*References*

Abraham, Alexandre, Fabian Pedregosa, Michael Eickenberg, Philippe Gervais, Andreas Mueller, Jean Kossaifi, Alexandre Gramfort, Bertrand Thirion, and Gael Varoquaux. 2014. "Machine Learning for Neuroimaging with Scikit-Learn." *Frontiers in Neuroinformatics* 8. https://doi.org/10.3389/fninf.2014.00014.

Andersson, Jesper L. R., Stefan Skare, and John Ashburner. 2003. "How to Correct Susceptibility Distortions in Spin-Echo Echo-Planar Images: Application to Diffusion Tensor Imaging." *NeuroImage* 20 (2): 870–88. https://doi.org/10.1016/S1053-8119(03)00336-7.

Avants, B. B., C. L. Epstein, M. Grossman, and J. C. Gee. 2008. “Symmetric Diffeomorphic Image Registration with Cross-Correlation: Evaluating Automated Labeling of Elderly and Neurodegenerative Brain.” *Medical Image Analysis* 12 (1): 26–41. https://doi.org/10.1016/j.media.2007.06.004.

Behzadi, Yashar, Khaled Restom, Joy Liau, and Thomas T. Liu. 2007. “A Component Based Noise Correction Method (CompCor) for BOLD and Perfusion Based fMRI.” *NeuroImage* 37 (1): 90–101. https://doi.org/10.1016/j.neuroimage.2007.04.042.

Ciric, R., William H. Thompson, R. Lorenz, M. Goncalves, E. MacNicol, C. J. Markiewicz, Y. O. Halchenko, et al. 2022. “TemplateFlow: FAIR-Sharing of Multi-Scale, Multi-Species Brain Models.” *Nature Methods* 19: 1568–71. https://doi.org/10.1038/s41592-022-01681-2.

Cox, Robert W., and James S. Hyde. 1997. “Software Tools for Analysis and Visualization of fMRI Data.” *NMR in Biomedicine* 10 (4-5): 171–78. https://doi.org/10.1002/(SICI)1099-1492(199706/08)10:4/5<171::AID-NBM453>3.0.CO;2-L.

Dale, Anders M., Bruce Fischl, and Martin I. Sereno. 1999. “Cortical Surface-Based Analysis: I. Segmentation and Surface Reconstruction.” *NeuroImage* 9 (2): 179–94. https://doi.org/10.1006/nimg.1998.0395.

Esteban, Oscar, Ross Blair, Christopher J. Markiewicz, Shoshana L. Berleant, Craig Moodie, Feilong Ma, Ayse Ilkay Isik, et al. 2018. “fMRIPrep 23.1.3.” *Software*. https://doi.org/10.5281/zenodo.852659.

Esteban, Oscar, Christopher Markiewicz, Ross W Blair, Craig Moodie, Ayse Ilkay Isik, Asier Erramuzpe Aliaga, James Kent, et al. 2019. “fMRIPrep: A Robust Preprocessing Pipeline for Functional MRI.” *Nature Methods* 16: 111–16. https://doi.org/10.1038/s41592-018-0235-4.

Evans, AC, AL Janke, DL Collins, and S Baillet. 2012. “Brain Templates and Atlases.” *NeuroImage* 62 (2): 911–22. https://doi.org/10.1016/j.neuroimage.2012.01.024.

Fonov, VS, AC Evans, RC McKinstry, CR Almli, and DL Collins. 2009. “Unbiased Nonlinear Average Age-Appropriate Brain Templates from Birth to Adulthood.” *NeuroImage* 47, Supplement 1: S102. https://doi.org/10.1016/S1053-8119(09)70884-5.

Glasser, Matthew F., Stamatios N. Sotiropoulos, J. Anthony Wilson, Timothy S. Coalson, Bruce Fischl, Jesper L. Andersson, Junqian Xu, et al. 2013. “The Minimal Preprocessing Pipelines for the Human Connectome Project.” *NeuroImage*, Mapping the connectome, 80: 105–24. https://doi.org/10.1016/j.neuroimage.2013.04.127.

Gorgolewski, K., C. D. Burns, C. Madison, D. Clark, Y. O. Halchenko, M. L. Waskom, and S. Ghosh. 2011. “Nipype: A Flexible, Lightweight and Extensible Neuroimaging Data Processing Framework in Python.” *Frontiers in Neuroinformatics* 5: 13. https://doi.org/10.3389/fninf.2011.00013.

Gorgolewski, Krzysztof J., Oscar Esteban, Christopher J. Markiewicz, Erik Ziegler, David Gage Ellis, Michael Philipp Notter, Dorota Jarecka, et al. 2018. “Nipype.” *Software*. https://doi.org/10.5281/zenodo.596855.

Greve, Douglas N, and Bruce Fischl. 2009. “Accurate and Robust Brain Image Alignment Using Boundary-Based Registration.” *NeuroImage* 48 (1): 63–72. https://doi.org/10.1016/j.neuroimage.2009.06.060.

Jenkinson, Mark, Peter Bannister, Michael Brady, and Stephen Smith. 2002. “Improved Optimization for the Robust and Accurate Linear Registration and Motion Correction of Brain Images.” *NeuroImage* 17 (2): 825–41. https://doi.org/10.1006/nimg.2002.1132.

Klein, Arno, Satrajit S. Ghosh, Forrest S. Bao, Joachim Giard, Yrjö Häme, Eliezer Stavsky, Noah Lee, et al. 2017. “Mindboggling Morphometry of Human Brains.” *PLOS Computational Biology* 13 (2): e1005350. https://doi.org/10.1371/journal.pcbi.1005350.

Lanczos, C. 1964. “Evaluation of Noisy Data.” *Journal of the Society for Industrial and Applied Mathematics Series B Numerical Analysis* 1 (1): 76–85. https://doi.org/10.1137/0701007.

Patriat, Rémi, Richard C. Reynolds, and Rasmus M. Birn. 2017. “An Improved Model of Motion-Related Signal Changes in fMRI.” *NeuroImage* 144, Part A (January): 74–82. https://doi.org/10.1016/j.neuroimage.2016.08.051.

Posse, Stefan, Stefan Wiese, Daniel Gembris, Klaus Mathiak, Christoph Kessler, Maria-Liisa Grosse-Ruyken, Barbara Elghahwagi, Todd Richards, Stephen R. Dager, and Valerij G. Kiselev. 1999. “Enhancement of BOLD-Contrast Sensitivity by Single-Shot Multi-Echo Functional MR Imaging.” *Magnetic Resonance in Medicine* 42 (1): 87–97. https://doi.org/10.1002/(SICI)1522-2594(199907)42:1<87::AID-MRM13>3.0.CO;2-O.

Power, Jonathan D., Anish Mitra, Timothy O. Laumann, Abraham Z. Snyder, Bradley L. Schlaggar, and Steven E. Petersen. 2014. “Methods to Detect, Characterize, and Remove Motion Artifact in Resting State fMRI.” *NeuroImage* 84 (Supplement C): 320–41. https://doi.org/10.1016/j.neuroimage.2013.08.048.

Reuter, Martin, Herminia Diana Rosas, and Bruce Fischl. 2010. “Highly Accurate Inverse Consistent Registration: A Robust Approach.” *NeuroImage* 53 (4): 1181–96. https://doi.org/10.1016/j.neuroimage.2010.07.020.

Satterthwaite, Theodore D., Mark A. Elliott, Raphael T. Gerraty, Kosha Ruparel, James Loughead, Monica E. Calkins, Simon B. Eickhoff, et al. 2013. “An improved framework for confound regression and filtering for control of motion artifact in the preprocessing of resting-state functional connectivity data.” *NeuroImage* 64 (1): 240–56. https://doi.org/10.1016/j.neuroimage.2012.08.052.

Tustison, N. J., B. B. Avants, P. A. Cook, Y. Zheng, A. Egan, P. A. Yushkevich, and J. C. Gee. 2010. “N4ITK: Improved N3 Bias Correction.” *IEEE Transactions on Medical Imaging* 29 (6): 1310–20. https://doi.org/10.1109/TMI.2010.2046908.

Zhang, Y., M. Brady, and S. Smith. 2001. “Segmentation of Brain MR Images Through a Hidden Markov Random Field Model and the Expectation-Maximization Algorithm.” *IEEE Transactions on Medical Imaging* 20 (1): 45–57. https://doi.org/10.1109/42.906424.

***Multi-echo ICA***

The fMRIPrep-preprocessed multi-echo data were further denoised using TE-dependent multi-echo ICA (tedana.py, version 23.0.1;[143–146]). To separate T2*-dependent BOLD-like signals from S0-dependent non-BOLD noise, PCA was performed using the Minimum Description Length (MDL) algorithm, which extracts the minimal number of components, followed by ICA. Tedana's automatic classification was manually verified. Noise components were removed, and the remaining components were recombined to generate the optimally denoised time series.

Finally, the denoised data were spatially normalized to MNI space using antsApplyTransforms (ANTs) and anatomical derivatives from fMRIPrep and spatially smoothed using a Gaussian kernel with a 6mm full width at half-maximum (FWHM).

#### *Functional Network Connectivity analysis*

We analyzed functional brain connectivity during the movie-viewing task using NeuroMark framework applied to the 30-minute fMRI time-series data per condition. We used the latest NeuroMark 2.2 multi-scale ICN templates[62,63]; spatial maps for the 14 subdomains across the 105 ICN templates are shown in **fig. S3**[63,120]. These ICN maps are standardized network expression strength at each voxel relative to the rest of the brain and are not mutually exclusive. For visualization, we followed prior work[120] by thresholding each ICN map at 1.96 and summed the maps within each subdomain.

Following Bajracharya et al., 2024, we applied Multi-Objective Optimization ICA with Reference (MOO-ICAR)[61,147] to estimate participant-specific ICN spatial maps and associated time courses. MOO-ICAR was implemented using the Group ICA of FMRI Toolbox (GIFT) package v4.0.6.5 (https://trendscenter.org/software/gift/). This approach incorporates group-level priors while preserving individual variability, yielding 105 individual-specific ICNs per run. As a result, for each participant and condition, we obtained ICN time series and spatial maps aligned with the NeuroMark 2.2 template across three movie viewing runs.

Before computing static functional network connectivity (sFNC), the ICN time series were preprocessed to minimize physiological and motion-related noise[61]: (1) linear, quadratic, and cubic detrending; (2) multiple regression of six motion realignment parameters and their temporal derivatives; (3) de-spiking to remove outliers; and (4) band-pass filtering (0.01-0.15 Hz). Preprocessed time series were then concatenated across the three runs for each condition, and sFNC matrices were computed as Pearson correlations between all ICN pairs, followed by Fisher $z$ transformation to improve normality for group-level tests. Each participant thus had two $105 \times 105$ FNC matrices (one per condition). To compare FNC between VT and VC, we performed paired t-tests on the Fisher $z$-transformed connectivity values for each ICN pair (VT−VC) across participants and applied the Bonferroni correction for multiple comparisons across 5,460 unique ICN pairs.

#### *Neurosynth functional decoding*

To facilitate systems-level analysis beyond original NeuroMark categorization, we regrouped the ICNs into a smaller set of functionally interpretable systems. This reorganization was guided by the NeuroMark subdomains, FNC condition sensitivity, and functional annotations from Neurosynth, allowing us to define eight major systems (**fig. S5-6**). These sytesms more coherently reflected the connectivity patterns observed in the FNC contrast, obtained from movie-viewing data.

To further characterize the three VT-related systems reorganized from NeuroMarks's triple-network domain, we used Neurosynth topic association maps, which highlight voxels preferentially associated with studies related to specific psychological topics. These topic maps were drawn from the "vs-topics-100" set (https://neurosynth.org/analyses/topics/v4-topics-50/), derived using Latent Dirichlet Allocation (LDA) from abstracts of all 11,406 articles in the Neurosynth database as of July, 2015, after excluding non-psychological topics[148]. We then calculated spatial correlation between each of eight systems (sum of ICN maps thresholded at 1.96) and the Neurosynth topic maps (**fig. S6**).

#### *Clustering analysis*

We applied *k*-means clustering to the FNC contrasts (VT−VC) for the 659 Bonferroni-significant ICN pairs. For each participant, we formed a 650-element vector of contrast values and then concatenated these vectors across participants. The optimal number of clusters (*k*) was selected from 5 to 15 using a permutation-based evaluation of silhouette values. For each k, the observed silhouette score was compared against a null distribution generated by randomly permuting the data, and the k with the highest Z-scored silhouette value was chosen, resulting in 12 clusters (**fig. S7**, **table S11**). We then ran *k*-means clustering with $k = 12$. Cluster centroids were calculated using correlation distance, defined as the component-wise mean of the points within each cluster after centering and normalizing each point to zero mean and unit standard deviation.

Next, we examined whether expression of each connectivity cluster was associated with cytokine responses. For each cytokine, we modeled the fold-change contrast between conditions (ΔFC = VT−VC) as a function of the corresponding cluster centroid scores, controlling for sex, age, and randomized condition order. The model formula was specified as: ΔFC ~ Cluster centroid + Condition Order + Sex + Age. Models were estimated using the restricted maximum likelihood (REML) option in fitlme function of Matlab.

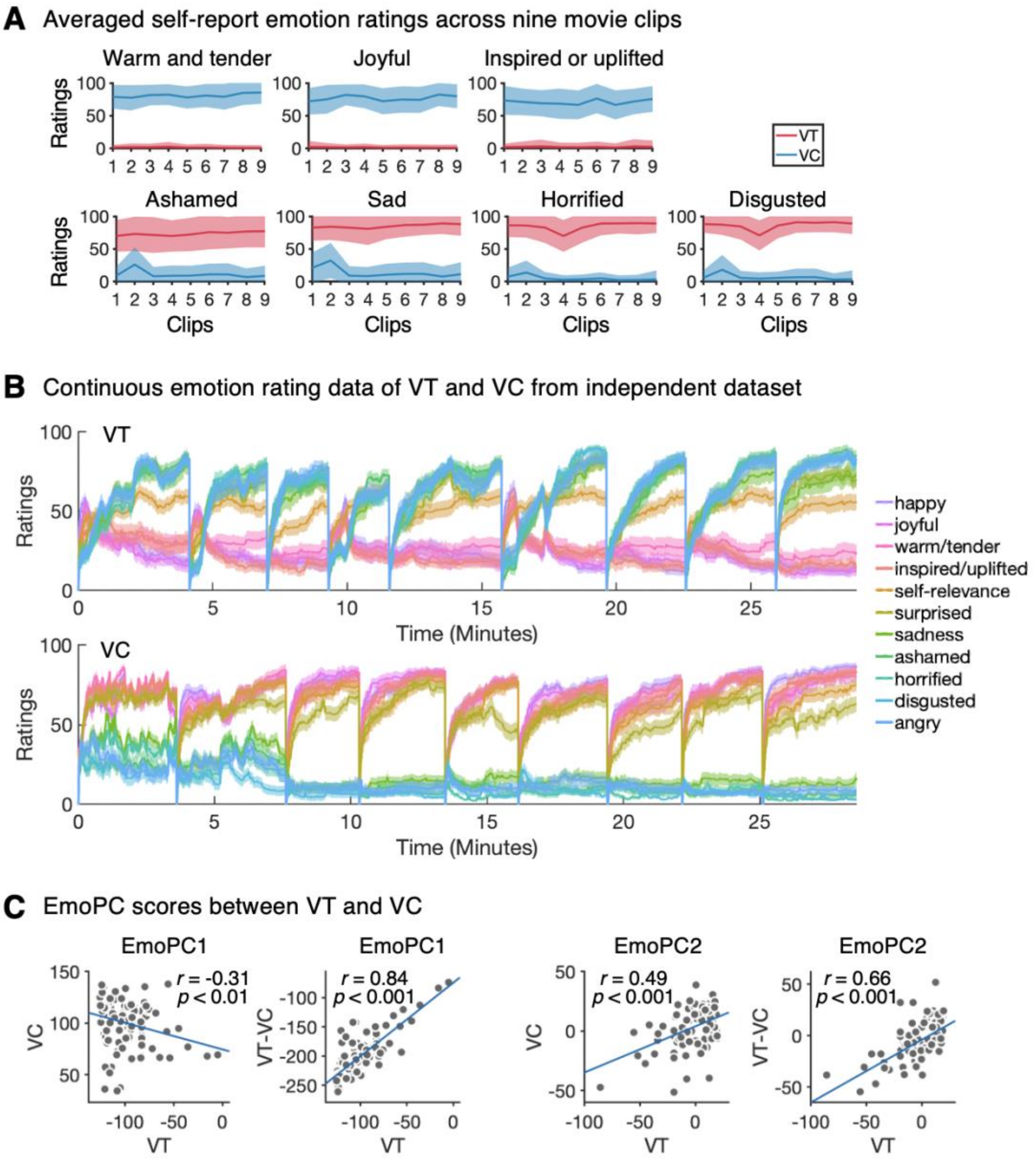


**Fig. S1. Self-report emotion ratings respond to movie clips**

(**A**) Mean self-report emotion ratings across movie clips. Lines show mean ratings for each condition, and shades indicate ±1 standard deviation. (**B**) Continuous emotion ratings for two films were obtained from an independent dataset sampling a broader range of emotions (total samples: VT, $n = 460$, VC, $n = 448$). In this online dataset, participants continuously rated a single target emotion while watching one of two videos ($n = 31$-57 per emotion). Ratings reliably distinguished the clips: consistent with the average ratings in the fMRI sample, VT clips evoked high negative and low positive affect, whereas VC clips produced the opposite pattern. (**C**) Average EmoPC1 and EmoPC2 scores by condition. EmoPC1 scores are participant-by-condition averages. EmoPC1 and EmoPC2 showed significant correlations across conditions (robust regression, $r = -0.31$, $P < 0.01$ and $r = 0.49$, $P < 0.001$). Participants' mean EmoPC scores in VT are correlated with their within-subject VT−VC differences (ΔEmoPC1, ΔEmoPC2).

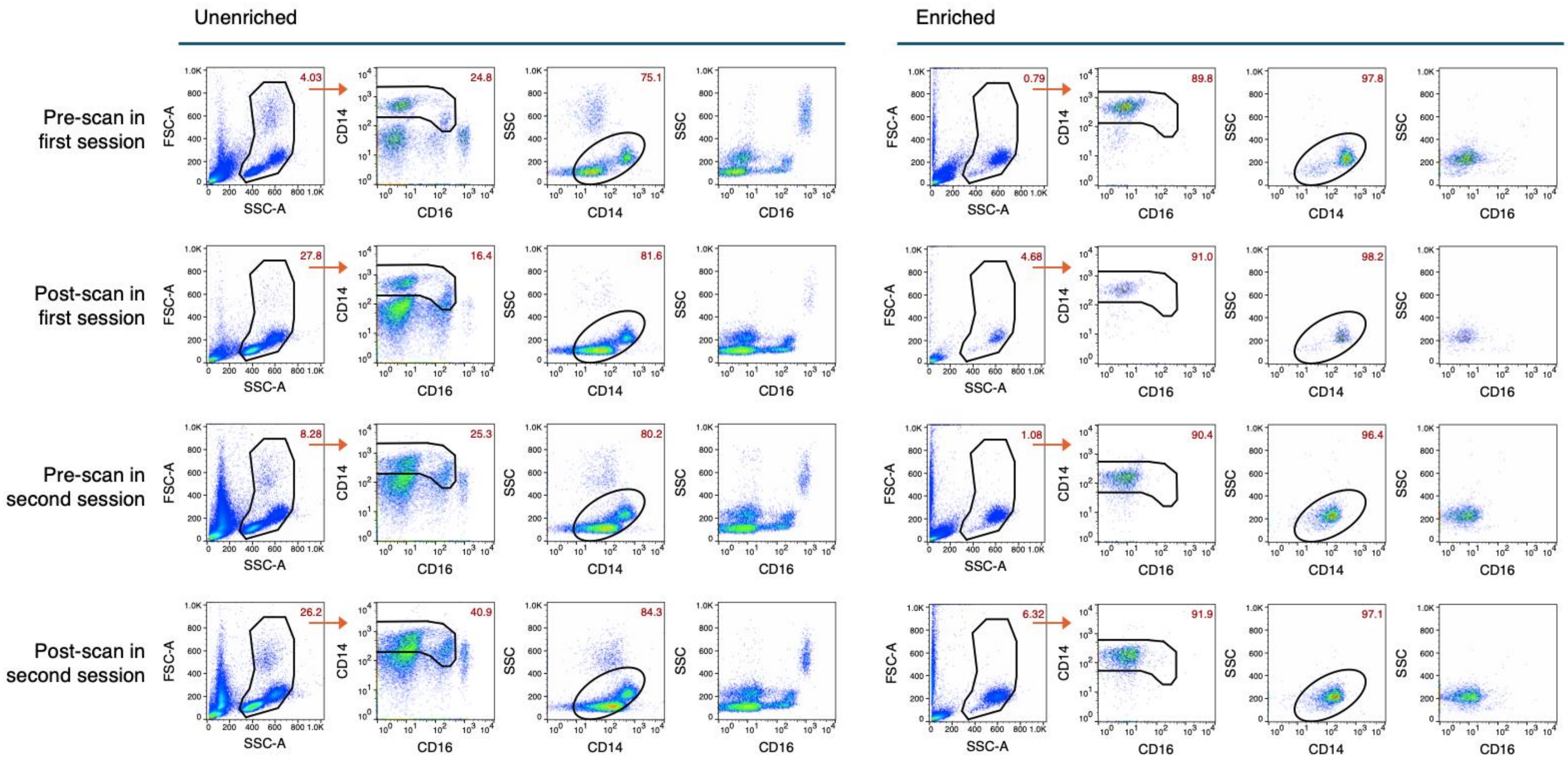


**Fig. S2. Representative flow cytometry results of monocyte enrichment**

Illustration of representative samples from a single participant (same participant as **Fig. 3A**) across two sessions. Rows (top to bottom) correspond to: pre-scan (First session), post-scan (Second session), pre-scan (Second session), and post-scan (Second session). Each row contains eight panels: the left four columns show enriched samples and the right four columns show enriched samples. For each sample, panels (left to right) are: dead cell exclusion and singlets were first performed, then analyzed using FSC vs SSC to gate on live cells (indicated by a black outline). Then we examined the live cell population by CD14 vs CD16 (gating on monocyte indicated by a black outline), SSC vs CD16, SSC vs CD14, (monocyte indicated by an oval outline). SSC vs CD16 and SSC vs CD14 distinguish intermediate-SSC monocytes from high-SSC CD16$^{high}$ neutrophils and low-SSC lymphocytes. Gated percentages are shown in the upper-right of each plot. The gated monocyte fraction increased from 16.4-40.9% pre-enrichment to 89.8-91.9% post-enrichment, confirming successful monocyte enrichment.

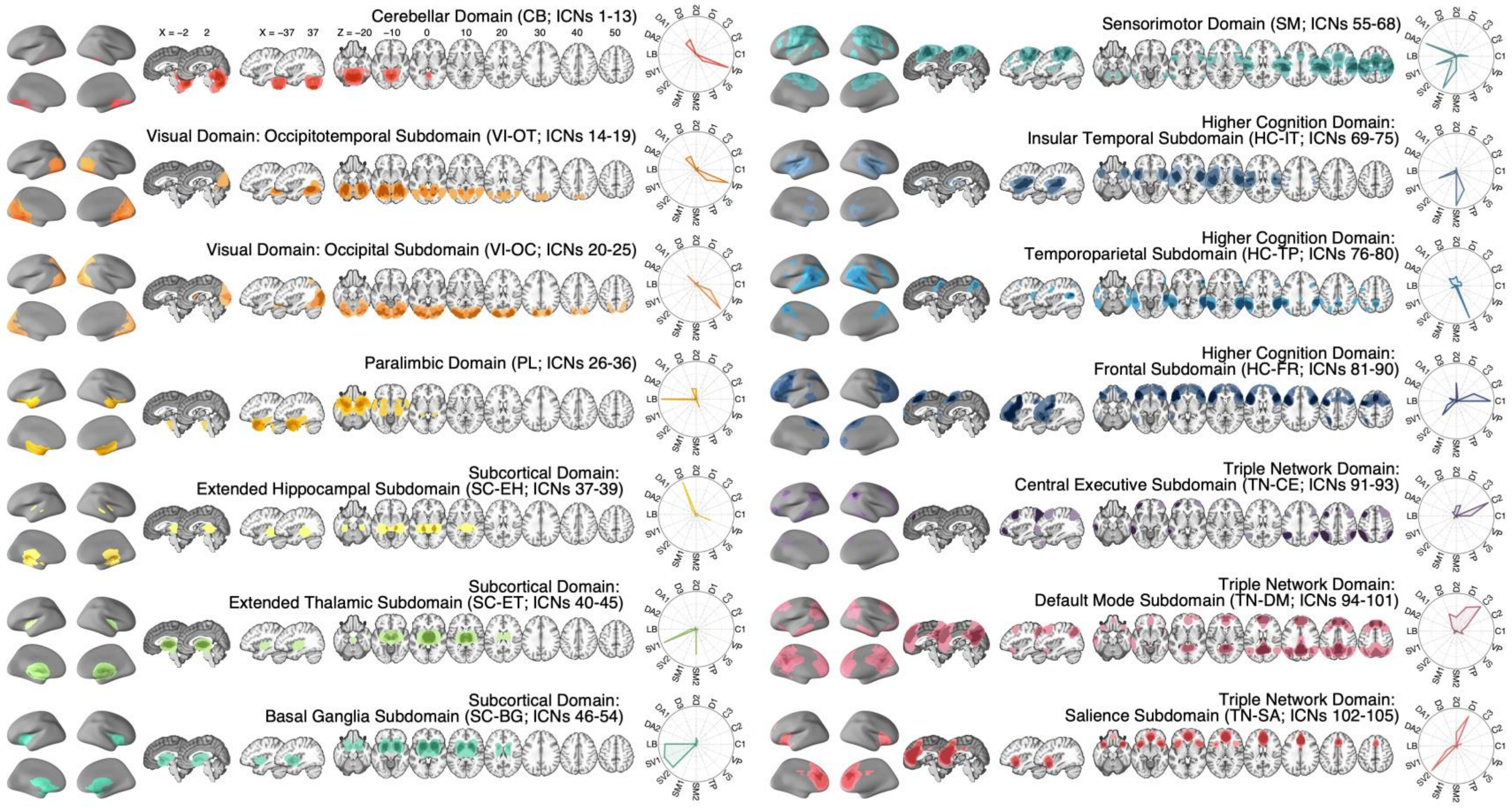


**Fig. S3. Visualization of 105 intrinsic connectivity networks in NeuroMark**

The NeuroMark 2.2 template includes 105 intrinsic connectivity networks (ICNs) categorized into 7 domains, four of which are further divided into subdomains, resulting in total of 14 subdomains[62,63,120]. Each subdomain was visualized with brain images with the ICN numbers. The map was generated by summing individual ICN masks that included voxels exceeding a threshold of 1.96, as provided in the original study. Darker regions therefore indicate area with greater overlap across ICNs within a given subdomain. The polar plots show the degree (%) of overlap with the 17 network parcellation from Schaefer et al., 2018[87].

Abbreviations: D1-D3 = Default A-C; C1-C3 = Control A-C; VP = Visual A; VS = Visual B; TP = Temporal-Parietal; SM1-2 = Somatomotor A-B; SV1-2 = Salience/Ventral Attention A-B; LB = Limbic.

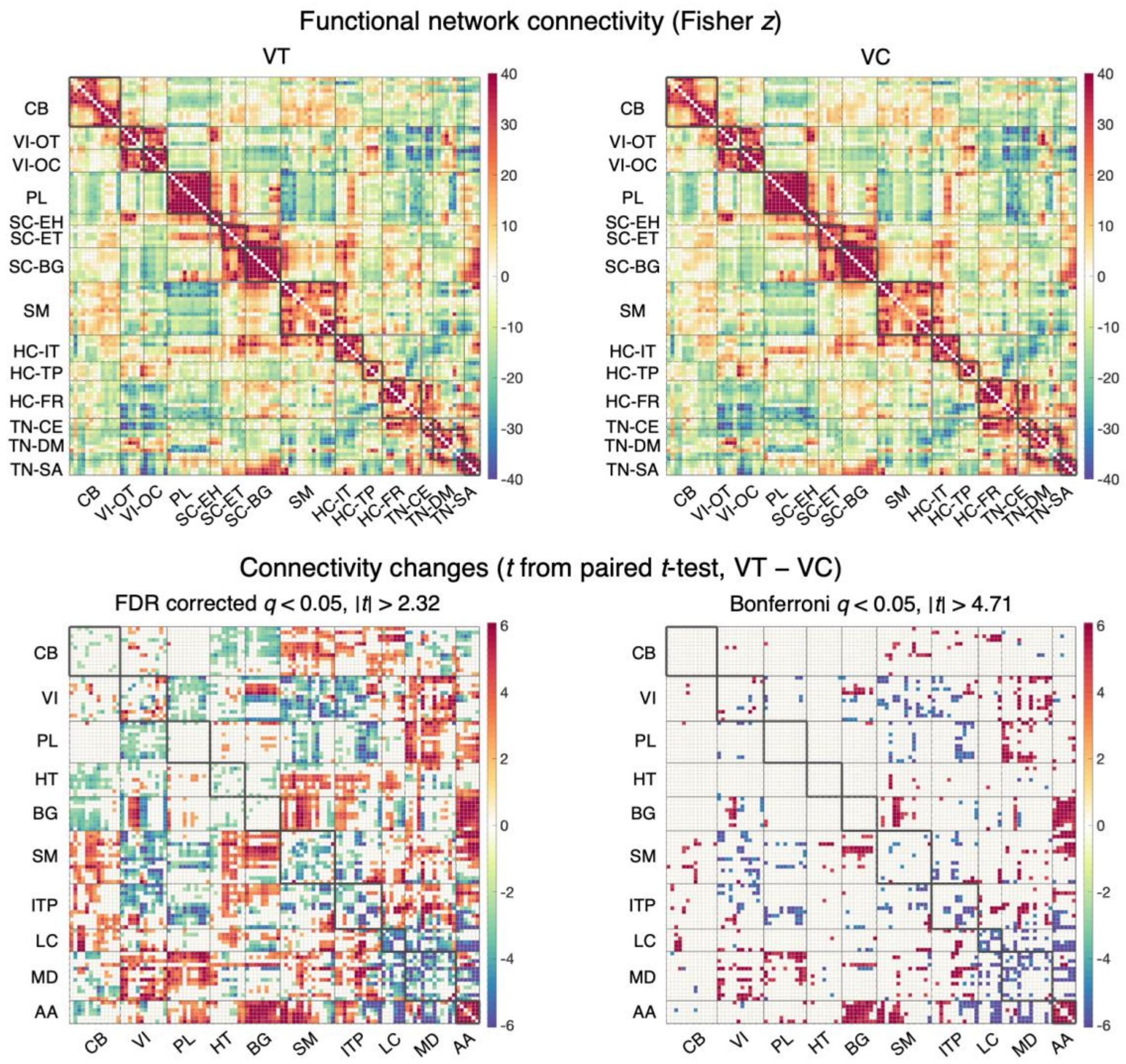


**Fig. S4. Functional network connectivity in VT and VC**

The top panel shows averaged correlation matrices of functional network connectivity (FNC; Fisher $z$) across 105 ICNs during movie watching in VT and VC ($n = 88$). The bottom panel shows connectivity differences ($t$ values) from paired $t$-tests (VT−VC), revealing significant condition-related changes across multiple networks. Among the 5,460 ICN pairs (105 × 104 / 2), 2,470 passed FDR correction at $q < 0.05$ ($|t| > 2.32$), and 659 passed Bonferroni correction at $q < 0.05$ ($|t| > 4.71$, same figure in **Fig. 4B**). Gray lines within the matrices indicate the boundaries of the Neuromark subdomains (top panel) and reorganized systems for this study (bottom panel). NeuroMark domain abbreviation are listed in **fig. S3**.

Abbreviations: CB, cerebellar; VI, visual; PL, paralimbic; HT, hippocampal-thalamic; BG, basal ganglia; SM, sensorimotor; ITP, insular-temporoparietal; LC, language-control; MD, memory-default; AA, affective-action.

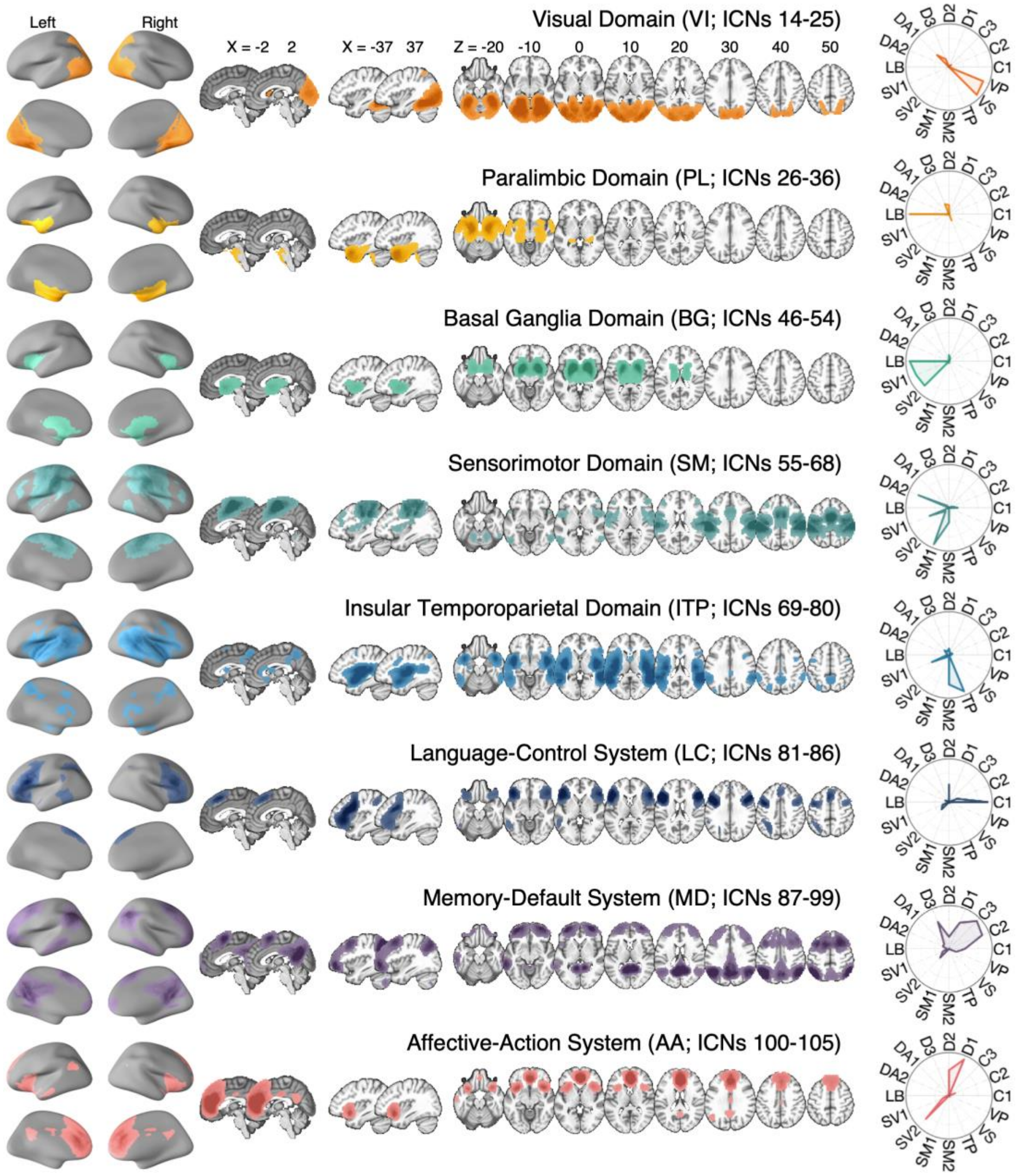


**Fig. S5. Visualization of eight major systems showing significant connectivity changes**
The NeuroMark template was originally developed from resting-state fMRI data, but here we applied it to movie-viewing fMRI. Additionally, the pattern of FNC difference across networks motivated a reorganization of the template to improve interpretability, resulting in eight major systems (**Fig. 4C**). This figure illustrates these systems, including their ICN numbers, brain maps, and polar plots indicating overlap percentage with the 17-network parcellation from Schaefer et al., 2018[87]. The labeling of the last three VT-related networks (LC, MD, and AA) was informed by Neurosynth topic mapping analysis (**fig. S6**).
Abbreviations: D1-D3 = Default A-C; C1-C3 = Control A-C; VP = Visual A; VS = Visual B; TP = Temporal-Parietal; SM1-2 = Somatomotor A-B; SV1-2 = Salience/Ventral Attention A-B; LB = Limbic.

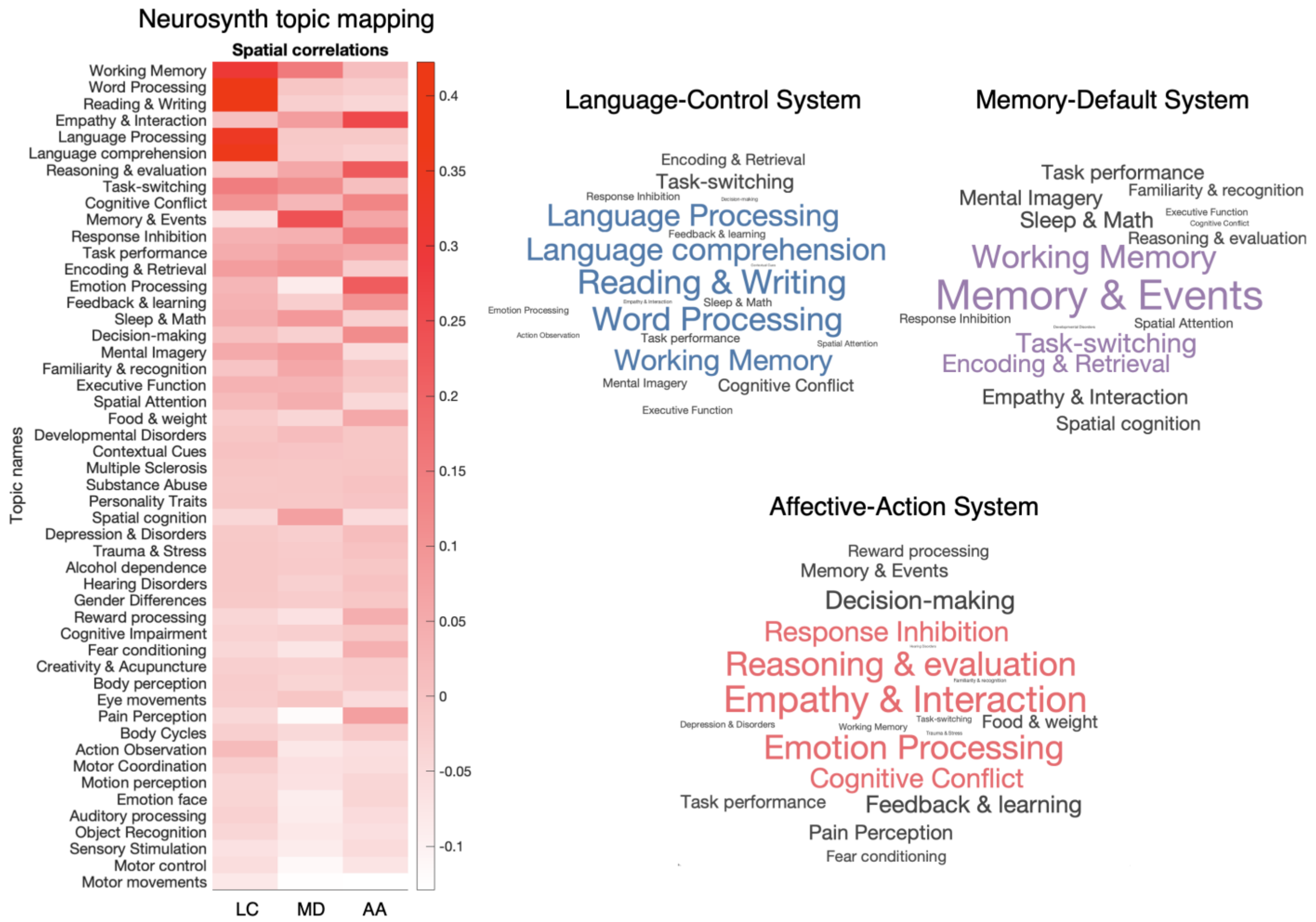


**Fig. S6. Neurosynth-based functional decoding of the three VT-related systems**

To functionally interpret the three VT-related systems (**Fig. 4**), we computed their associations with 50 Neurosynth psychological topic maps related to mental process[86] (https://neurosynth.org/; reverse inference maps). The left panel shows the spatial correlations between each system and the topic maps, sorted by the summed correlations across the three systems. The right panel shows word clouds highlighting the most relevant topics for each system, which guided our labeling of the systems as Language-Control (LC), Memory-Default (MD), and Affective-Action (AA) systems.

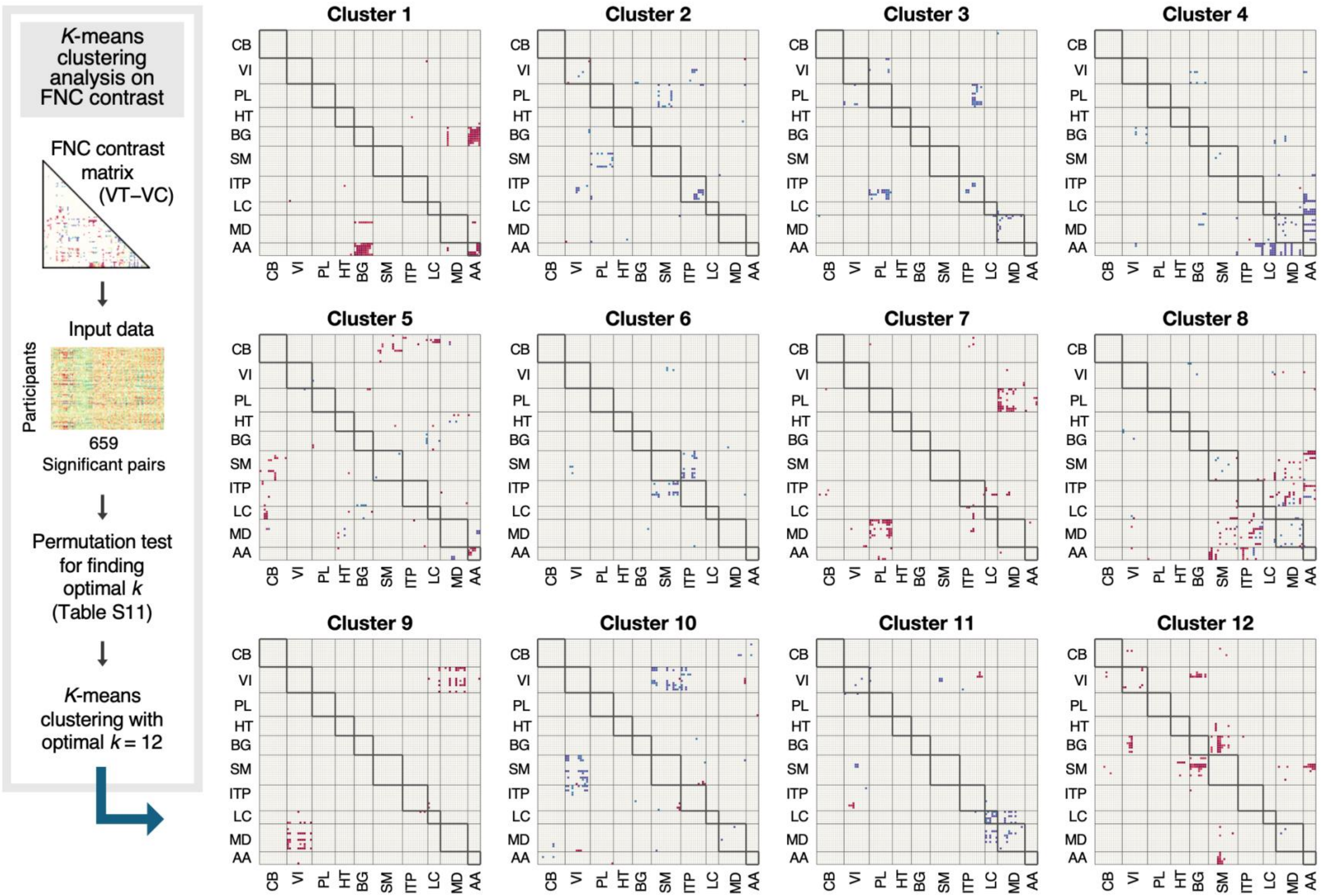


**Fig. S7. Clustering results of functional network connectivity**

Illustration of the 12 clusters derived from *k*-means clustering of the 659 Bonferroni-significant ICN pairs. Each matrix shows the FNC contrast (VT−VC in **Fig. 4B**) for the corresponding ICN belonging to that cluster. Red indicates VT > VC, while blue indicates VT < VC connectivity.
Abbreviations: CB, cerebellar; VI, visual; PL, paralimbic; HT, hippocampal-thalamic; BG, basal ganglia; SM, sensorimotor; ITP, insular-temporoparietal; LC, language-control; MD, memory-default; AA, affective-action.

**Table S1. Descriptive statistics of self-reported emotion ratings**

| | Vicarious Trauma | | Vicarious Community | |
|---|---|---|---|---|
| Emotions | Mean | STD | Mean | STD |
| Warm and tender | 1.66 | 4.21 | 80.97 | 14.03 |
| Joyful | 1.68 | 4.10 | 76.94 | 14.90 |
| Inspired or Uplifted | 2.50 | 6.73 | 71.05 | 16.70 |
| Ashamed | 73.57 | 21.49 | 11.14 | 11.35 |
| Sad | 85.12 | 14.19 | 13.89 | 12.59 |
| Horrified | 85.03 | 15.49 | 5.27 | 6.85 |
| Disgusted | 86.51 | 13.74 | 6.82 | 7.69 |

Mean and standard deviation of seven emotion ratings for nine movie clips for two conditions ($n = 88$)

**Table S2. Autonomic responses paired *t*-test results (VT−VC)**

| | Heart rate (BPM) | | | | Skin conductance | | | | Heart rate variability | | | |
|---|---|---|---|---|---|---|---|---|---|---|---|---|
| | *t* | | *P* | *DF* | *t* | | *P* | *DF* | *t* | | *P* | *DF* |
| Pre | 1.62 | | 0.110 | 84 | 1.28 | | 0.206 | 71 | -0.87 | | 0.387 | 84 |
| Movie | 2.40 | * | 0.019 | 84 | 3.95 | *** | 0.0002 | 71 | 1.42 | | 0.160 | 84 |
| Rest | 2.98 | ** | 0.004 | 84 | 2.51 | * | 0.015 | 71 | -2.73 | ** | 0.008 | 84 |
| Post-scan | 2.41 | * | 0.018 | 84 | 0.28 | | 0.779 | 71 | -0.52 | | 0.606 | 84 |

Paired *t*-test results for autonomic responses across the experiment (VT−VC). Pre: The 20 seconds before the movie began (−20 to 0 seconds); Movie: the 30-minute movie-viewing fMRI runs; Rest: the 10-minute resting-state fMRI run; Post-scan: the 5-minute resting autonomic recording. Participants already knew which video they would watch during the Pre period, which may have contribute to differences in that interval, although the Pre differences were not significant. *$P < .05$. **$P < .01$. ***$P < .001$.

**Table S3. Linear mixed-effects modeling predicting heart rate during movie watching from EmoPCs**

| Fixed effects | Estimate | SE | *t* | *df* | *P* | | Lower | Upper |
|---|---|---|---|---|---|---|---|---|
| Intercept | 63.6840 | 0.9452 | 67.37 | 84.0 | 6.97E-75 | | 61.8040 | 65.5630 |
| EmoPC1 | -0.0099 | 0.0014 | -7.31 | 1443.2 | 4.26E-13 | | -0.0126 | -0.0073 |
| EmoPC2 | 0.0062 | 0.0078 | 0.79 | 1481.9 | 0.4320 | | -0.0092 | 0.0215 |
| PC1:PC2 | -0.0003 | 0.0001 | -3.36 | 1455.3 | 0.0008 | *** | -0.0005 | -0.0001 |

| Random effects | Group | Effect | Estimate |
|---|---|---|---|
| | Subject | (Intercept) Std. Dev. | 0.303 |
| | Subject | Correlation: Cond & Intercept | -0.353 |
| | Subject | Cond Std. Dev. | 0.272 |
| | Error | Residual Std. Dev. | 0.278 |

Linear mixed-effects model of heart rate during movie watching predicted by EmoPC scores ($n = 85$). The model formula was heart rate ~ EmoPC1 * EmoPC2 + (1 + Clip | Subject), with participants modeled as random intercepts and clip numbers as random slopes. A significant EmoPC1 × EmoPC2 interaction indicated that higher heart rate was associated with more negative valence (lower EmoPC1), especially for clips where participants reported greater empathic engagement (higher EmoPC2) (**Fig. 2C**). *$P < .05$. **$P < .01$. ***$P < .001$.

**Table S4. Within-subject reliability of baseline gene expression for target cytokines**

| Gene | IL1B | CXCL8 | TNF | TGFB1 |
|---|---|---|---|---|
| r | 0.355 | 0.253 | 0.274 | 0.347 |
| *P* | 0.008 ** | 0.063 | 0.043 * | 0.010 * |

Three of four target cytokines showed significant baseline correlations across the two conditions ($n$ = 55). Although CXCL8 did not reach significance, the correlation was positive.
*$P$ < .05. **$P$ < .01.

**Table S5. One-sample and paired *t*-test results for fold changes in target cytokines**

| Gene | IL1B | | | CXCL8 | | | TNF | | | TGFB1 | | |
|---|---|---|---|---|---|---|---|---|---|---|---|---|
| Condition | VT | VC | VT−VC | VT | VC | VT−VC | VT | VC | VT−VC | VT | VC | VT−VC |
| Mean | -0.188 | -0.284 | 0.096 | -0.604 | -0.957 | 0.353 | 0.075 | 0.038 | 0.037 | -0.212 | 0.002 | -0.213 |
| *t* | -1.749 | -2.861 | 0.805 | -2.824 | -5.790 | 1.490 | 1.064 | 0.688 | 0.459 | -2.792 | 0.033 | -2.502 |
| *P* | 0.086 | 0.006 ** | 0.424 | 0.007 ** | 0.000 *** | 0.142 | 0.292 | 0.495 | 0.648 | 0.007 ** | 0.974 | 0.015 * |
| Cohen's *d* | -0.236 | -0.386 | 0.109 | -0.381 | -0.781 | 0.201 | 0.143 | 0.093 | 0.062 | -0.377 | 0.004 | -0.337 |

One sample *t*-tests assessed fold changes in cytokines within each condition, and paired *t*-tests compared fold changes between conditions (VT−VC) ($n$ = 55). IL1B showed a significant reduction after VC. CXCL8 showed significant reductions after both conditions. TGFB1 showed a significant reduction after VT, and it was the only cytokine that showed a greater reduction after VT compared to VC. *$P < .05$. **$P < .01$. ***$P < .001$.

**Table S6. Condition-level linear mixed-effects modeling predicting post-scan cytokine gene expression**

IL1B

| Fixed effects | Estimate | *SE* | *t* | *DF* | *P* | | Lower | Upper |
|---|---|---|---|---|---|---|---|---|
| Intercept | 34.605 | 3.406 | 10.16 | 51.55 | 0.0000 | *** | 27.769 | 41.442 |
| Condition | -1.175 | 1.045 | -1.13 | 53.48 | 0.2656 | | -3.270 | 0.920 |
| Order | -0.826 | 0.794 | -1.04 | 49.83 | 0.3036 | | -2.421 | 0.770 |
| Sex | 0.063 | 0.793 | 0.08 | 49.76 | 0.9366 | | -1.530 | 1.657 |
| Age | -0.044 | 0.142 | -0.31 | 50.07 | 0.7600 | | -0.329 | 0.242 |
| Batch | -1.582 | 0.347 | -4.56 | 51.42 | 0.0000 | *** | -2.279 | -0.886 |
| Pre | 0.304 | 0.035 | 8.68 | 80.28 | 0.0000 | *** | 0.235 | 0.374 |
| Cond:Pre | -0.094 | 0.037 | -2.55 | 93.15 | 0.0125 | * | -0.168 | -0.021 |
| **Random effects** | **Type** | **Estimate** | | | | | **Lower** | **Upper** |
| Intercept | STD | 5.951 | | | | | NaN | NaN |
| Intercept & Condition | Corr | -0.682 | | | | | NaN | NaN |
| Condition | STD | 7.012 | | | | | NaN | NaN |
| Error (Residual) | STD | 4.647 | | | | | NaN | NaN |

CXCL8

| Fixed effects | Estimate | *SE* | *t* | *DF* | *P* | | Lower | Upper |
|---|---|---|---|---|---|---|---|---|
| Intercept | 10.176 | 2.880 | 3.53 | 51.80 | 0.0009 | *** | 4.397 | 15.955 |
| Condition | -1.063 | 0.775 | -1.37 | 52.28 | 0.1761 | | -2.618 | 0.492 |
| Order | 0.354 | 0.651 | 0.54 | 48.57 | 0.5891 | | -0.955 | 1.663 |
| Sex | 0.945 | 0.648 | 1.46 | 48.56 | 0.1510 | | -0.357 | 2.247 |
| Age | -0.071 | 0.117 | -0.61 | 49.04 | 0.5454 | | -0.307 | 0.164 |
| Batch | -0.476 | 0.310 | -1.54 | 51.56 | 0.1306 | | -1.097 | 0.146 |
| Pre | 0.271 | 0.044 | 6.15 | 79.27 | 0.0000 | *** | 0.183 | 0.359 |
| Cond:Pre | -0.089 | 0.041 | -2.19 | 63.41 | 0.0324 | * | -0.170 | -0.008 |
| **Random effects** | **Type** | **Estimate** | | | | | **Lower** | **Upper** |
| Intercept | STD | 5.217 | | | | | NaN | NaN |
| Intercept & Condition | Corr | -0.771 | | | | | NaN | NaN |
| Condition | STD | 5.051 | | | | | NaN | NaN |
| Error (Residual) | STD | 3.883 | | | | | NaN | NaN |

TNF

| Fixed effects | Estimate | *SE* | *t* | *DF* | *P* | | Lower | Upper |
|---|---|---|---|---|---|---|---|---|
| Intercept | 13.125 | 1.423 | 9.23 | 48.51 | 0.0000 | *** | 10.266 | 15.985 |
| Condition | -0.038 | 0.238 | -0.16 | 51.44 | 0.8730 | | -0.516 | 0.440 |
| Order | -0.133 | 0.337 | -0.40 | 49.64 | 0.6943 | | -0.811 | 0.544 |
| Sex | 0.451 | 0.333 | 1.36 | 48.89 | 0.1817 | | -0.218 | 1.120 |
| Age | -0.052 | 0.060 | -0.86 | 49.14 | 0.3937 | | -0.173 | 0.069 |
| Batch | -0.142 | 0.153 | -0.93 | 53.31 | 0.3557 | | -0.449 | 0.164 |
| Pre | 0.594 | 0.074 | 8.05 | 101.76 | 0.0000 | *** | 0.448 | 0.740 |
| Cond:Pre | -0.015 | 0.061 | -0.25 | 70.57 | 0.8010 | | -0.137 | 0.106 |
| **Random effects** | **Type** | **Estimate** | | | | | **Lower** | **Upper** |
| Intercept | STD | 2.124 | | | | | NaN | NaN |
| Intercept & Condition | Corr | -0.191 | | | | | NaN | NaN |
| Condition | STD | 1.295 | | | | | NaN | NaN |
| Error (Residual) | STD | 1.697 | | | | | NaN | NaN |

TGFB1

| Fixed effects | Estimate | *SE* | *t* | *DF* | *P* | | Lower | Upper |
|---|---|---|---|---|---|---|---|---|
| Intercept | 373.830 | 71.215 | 5.25 | 49.03 | 0.0000 | *** | 230.720 | 516.940 |
| Condition | 30.864 | 13.629 | 2.26 | 51.43 | 0.0278 | * | 3.508 | 58.220 |
| Order | -16.442 | 16.358 | -1.01 | 47.60 | 0.3199 | | -49.340 | 16.456 |
| Sex | 6.879 | 16.438 | 0.42 | 48.22 | 0.6774 | | -26.167 | 39.926 |
| Age | 1.018 | 2.931 | 0.35 | 47.33 | 0.7298 | | -4.876 | 6.913 |
| Batch | 20.422 | 7.801 | 2.62 | 54.39 | 0.0114 | * | 4.784 | 36.059 |
| Pre | 0.304 | 0.053 | 5.77 | 89.01 | 0.0000 | *** | 0.200 | 0.409 |
| Cond:Pre | 0.054 | 0.044 | 1.23 | 66.29 | 0.2222 | | -0.033 | 0.142 |
| **Random effects** | **Type** | **Estimate** | | | | | **Lower** | **Upper** |
| Intercept | STD | 100.540 | | | | | NaN | NaN |
| Intercept & Condition | Corr | 0.055 | | | | | NaN | NaN |
| Condition | STD | 78.263 | | | | | NaN | NaN |
| Error (Residual) | STD | 89.346 | | | | | NaN | NaN |

Linear mixed-effects models were conducted to predict post-scan gene expression (CPM) using condition and baseline ($n$ = 55). The model formula was: Post ~ Condition * Pre (Baseline) + Condition Order + Sex + Age + Batch + (1 + Condition | Subject), with partic ipants modeled as random intercepts and condition as random slopes. Baseline expression significantly predicted post-scan levels for a ll four cytokines. IL1B and CXCL8 showed significant condition × baseline interactions (**Fig. 3F**). TGFB1 also showed a significant main effect of condition, consistent with the paired $t$-test results (Table S5). *$P$ < .05. **$P$ < .01. ***$P$ < .001.

**Table S7. Condition-level linear mixed-effects modeling predicting cytokine fold change from EmoPCs and heart rate**

IL1B

| Fixed effects | Estimate | *SE* | *t* | *DF* | *P* | | Lower | Upper |
|---|---|---|---|---|---|---|---|---|
| Intercept | -1.533 | 0.635 | -2.42 | 76.02 | 0.0181 | * | -2.797 | -0.269 |
| Order | 0.013 | 0.084 | 0.15 | 46.45 | 0.8809 | | -0.157 | 0.182 |
| Sex | 0.010 | 0.087 | 0.12 | 48.41 | 0.9082 | | -0.164 | 0.184 |
| Age | -0.009 | 0.015 | -0.59 | 46.63 | 0.5602 | | -0.039 | 0.022 |
| Heart Rate | 0.024 | 0.008 | 2.79 | 90.27 | 0.0064 | ** | 0.007 | 0.041 |
| EmoPC1 | 0.000 | 0.001 | -0.12 | 54.23 | 0.9019 | | -0.001 | 0.001 |
| EmoPC2 | -0.001 | 0.004 | -0.34 | 99.00 | 0.7337 | | -0.010 | 0.007 |
| Random effects | Type | Estimate | | | | | Lower | Upper |
| Intercept | STD | 0.421 | | | | | 0.264 | 0.673 |
| Error (Residual) | STD | 0.602 | | | | | 0.494 | 0.734 |

CXCL8

| Fixed effects | Estimate | *SE* | *t* | *DF* | *P* | | Lower | Upper |
|---|---|---|---|---|---|---|---|---|
| Intercept | -1.153 | 1.316 | -0.88 | 73.91 | 0.3840 | | -3.775 | 1.470 |
| Order | 0.062 | 0.170 | 0.37 | 48.84 | 0.7157 | | -0.280 | 0.404 |
| Sex | 0.037 | 0.175 | 0.21 | 50.66 | 0.8325 | | -0.315 | 0.389 |
| Age | -0.041 | 0.031 | -1.33 | 49.01 | 0.1902 | | -0.102 | 0.021 |
| Heart Rate | 0.019 | 0.018 | 1.07 | 85.52 | 0.2884 | | -0.016 | 0.054 |
| EmoPC1 | -0.002 | 0.001 | -1.30 | 56.87 | 0.1977 | | -0.004 | 0.001 |
| EmoPC2 | 0.010 | 0.009 | 1.09 | 97.86 | 0.2805 | | -0.008 | 0.027 |
| Random effects | Type | Estimate | | | | | Lower | Upper |
| Intercept | STD | 0.734 | | | | | 0.390 | 1.381 |
| Error (Residual) | STD | 1.359 | | | | | 1.121 | 1.648 |

TNF

| Fixed effects | Estimate | *SE* | *t* | *DF* | *P* | | Lower | Upper |
|---|---|---|---|---|---|---|---|---|
| Intercept | 0.185 | 0.396 | 0.47 | 63.52 | 0.6418 | | -0.607 | 0.977 |
| Order | -0.076 | 0.050 | -1.52 | 42.20 | 0.1363 | | -0.176 | 0.025 |
| Sex | 0.042 | 0.051 | 0.81 | 43.81 | 0.4200 | | -0.062 | 0.146 |
| Age | -0.008 | 0.009 | -0.89 | 42.35 | 0.3779 | | -0.026 | 0.010 |
| Heart Rate | 0.000 | 0.005 | 0.07 | 73.87 | 0.9476 | | -0.010 | 0.011 |
| EmoPC1 | 0.000 | 0.000 | -0.35 | 50.51 | 0.7300 | | -0.001 | 0.001 |
| EmoPC2 | 0.000 | 0.003 | -0.11 | 91.95 | 0.9107 | | -0.006 | 0.005 |
| Random effects | Type | Estimate | | | | | Lower | Upper |
| Intercept | STD | 0.146 | | | | | 0.026 | 0.812 |
| Error (Residual) | STD | 0.456 | | | | | 0.371 | 0.561 |

TGFB1

| Fixed effects | Estimate | *SE* | *t* | *DF* | *P* | | Lower | Upper |
|---|---|---|---|---|---|---|---|---|
| Intercept | 0.312 | 0.394 | 0.79 | 72.60 | 0.4298 | | -0.472 | 1.097 |
| Order | -0.060 | 0.050 | -1.19 | 48.69 | 0.2402 | | -0.162 | 0.041 |
| Sex | -0.051 | 0.052 | -0.97 | 50.47 | 0.3349 | | -0.155 | 0.054 |
| Age | -0.001 | 0.009 | -0.15 | 48.86 | 0.8795 | | -0.020 | 0.017 |
| Heart Rate | -0.006 | 0.005 | -1.15 | 83.62 | 0.2543 | | -0.017 | 0.004 |
| EmoPC1 | 0.001 | 0.000 | 2.45 | 56.81 | 0.0174 | * | 0.000 | 0.002 |
| EmoPC2 | -0.001 | 0.003 | -0.46 | 96.99 | 0.6480 | | -0.006 | 0.004 |
| Random effects | Type | Estimate | | | | | Lower | Upper |
| Intercept | STD | 0.203 | | | | | 0.097 | 0.427 |
| Error (Residual) | STD | 0.418 | | | | | 0.345 | 0.507 |

Linear mixed-effects models were conducted to predict cytokine fold changes using EmoPCs and heart rate during movie watching ($n$ = 55). The model formula was: Fold change ~ EmoPC1 + EmoPC2 + Heart rate + Condition Order + Sex + Age + (1 | Subject). The condition variable was excluded because it was highly correlated with EmoPC1. Heart rate during movie watching significantly predicted IL1B fold change. EmoPC1 significantly predicted TGFB1 fold change, consistent with the VT-related reduction.
*$P < .05$. **$P < .01$.

**Table S8. Condition-level linear mixed-effects modeling predicting *IL-1β* fold change from EmoPCs and heart rate with interactions**

| IL1B | | | | | | | | |
|---|---|---|---|---|---|---|---|---|
| Fixed effects | Estimate | *SE* | *t* | *DF* | *P* | | Lower | Upper |
| Intercept | -1.259 | 0.631 | -2.00 | 69.63 | 0.0499 | * | -2.517 | -0.001 |
| Order | 0.005 | 0.081 | 0.06 | 44.18 | 0.9500 | | -0.157 | 0.167 |
| Sex | 0.021 | 0.083 | 0.25 | 47.76 | 0.8028 | | -0.147 | 0.189 |
| Age | -0.008 | 0.015 | -0.58 | 44.55 | 0.5650 | | -0.038 | 0.021 |
| Heart Rate | 0.019 | 0.009 | 2.18 | 83.07 | 0.0320 | * | 0.002 | 0.036 |
| EmoPC1 | 0.008 | 0.004 | 1.96 | 62.05 | 0.0543 | | 0.000 | 0.017 |
| EmoPC2 | -0.001 | 0.004 | -0.21 | 96.68 | 0.8370 | | -0.009 | 0.007 |
| HR:EmoPC1 | 0.000 | 0.000 | -2.03 | 63.37 | 0.0467 | * | 0.000 | 0.000 |
| EmoPC1:PC2 | 0.000 | 0.000 | 0.54 | 71.11 | 0.5926 | | 0.000 | 0.000 |
| Random effects | Type | Estimate | | | | | Lower | Upper |
| Intercept | STD | 0.371 | | | | | 0.204 | 0.673 |
| Error (Residual) | STD | 0.616 | | | | | 0.502 | 0.757 |

Linear mixed-effects models were conducted to predict cytokine fold changes using EmoPCs and heart rate during movie watching including interactions ($n$ = 55). The model formula was: Fold change ~ EmoPC1 * EmoPC2 + EmoPC1 * Heart rate + Condition Order + Sex + Age + (1 | Subject). Heart rate during movie watching remained a significant predictor of IL1B fold change, and the heart rate × EmoPC1 interaction also significantly predicted IL1B fold change (**Fig. 3G**). *$P$ < .05.

**Table S9. Within-subject stability of spatial ICN maps across runs in two conditions**

| Condition | | VT | | VC | | |
|---|---|---|---|---|---|---|
| Run | | Run 2 | Run 3 | Run 1 | Run 2 | Run 3 |
| VT | Run 1 | 0.755 | 0.752 | 0.730 | 0.731 | 0.730 |
| | Run 2 | | 0.758 | 0.732 | 0.732 | 0.733 |
| | Run 3 | | | 0.732 | 0.733 | 0.734 |
| VC | Run 1 | | | | 0.755 | 0.752 |
| | Run 2 | | | | | 0.762 |

Averaged Fisher *z* scores are shown for within-subject correlations of spatial ICN maps across runs ($n$ = 88). Each condition included three runs. For each participant and each of the individualized ICN maps, spatial correlations across runs were computed and converted the Pearson's correlation to Fisher *z* scores. The *z* scores were then averaged across maps and participants for each run pair. The mean within-condition spatial correlation (VT & VT or VC & VC) was 0.756, and the mean cross-condition correlation (VT & VC) was 0.732.

**Table S10. Effect sizes of functional connectivity differences**

| Cohen's d | | | | | | | | |
|---|---|---|---|---|---|---|---|---|
| | VI | PL | BG | SM | ITP | LC | MD | AA |
| VI | 0.215 | | | | | | | |
| PL | -0.747 | NaN | | | | | | |
| BG | 0.105 | 0.532 | NaN | | | | | |
| SM | -0.600 | -0.585 | 0.582 | -0.301 | | | | |
| ITP | -0.375 | -0.718 | -0.346 | -0.374 | -0.707 | | | |
| LC | 0.501 | NaN | -0.276 | 0.657 | 0.580 | -0.649 | | |
| MD | 0.772 | 0.582 | 0.179 | 0.357 | 0.383 | -0.719 | -0.666 | |
| AA | 0.005 | 0.566 | 0.612 | 0.521 | -0.200 | -0.759 | -0.664 | 0.672 |

| *SD* | | | | | | | | |
|---|---|---|---|---|---|---|---|---|
| | VI | PL | BG | SM | ITP | LC | MD | AA |
| VI | 0.580 | | | | | | | |
| PL | 0.251 | NaN | | | | | | |
| BG | 0.578 | 0.015 | NaN | | | | | |
| SM | 0.245 | 0.071 | 0.068 | 0.486 | | | | |
| ITP | 0.542 | 0.240 | 0.439 | 0.552 | 0.427 | | | |
| LC | 0.420 | NaN | 0.614 | 0.080 | 0.428 | 0.134 | | |
| MD | 0.233 | 0.253 | 0.543 | 0.586 | 0.566 | 0.134 | 0.367 | |
| AA | 0.676 | 0.048 | 0.068 | 0.409 | 0.663 | 0.142 | 0.548 | 0.099 |

Among major eight systems, functional connectivity differences (VT−VC) were calculated, and the corresponding Cohen's d values and standard deviations were computed (**Fig. 4C**, $n$ = 88). Abbreviations for systems: VI, visual; PL, paralimbic; BG, basal ganglia; SM, sensorimotor; ITP, insular-temporoparietal; LC, language-control; MD, memory-default; AA, affective-action.

**Table S11. Permutation test results for determining the optimal number of clusters (*k*) for significant functional connectivity pairs**

| Cluster num | 5 | 6 | 7 | 8 | 9 | 10 | 11 | 12 | 13 | 14 | 15 |
|---|---|---|---|---|---|---|---|---|---|---|---|
| Quality | 0.079 | 0.082 | 0.080 | 0.083 | 0.084 | 0.083 | 0.088 | 0.092 | 0.092 | 0.092 | 0.096 |
| Null q *mean* | 0.004 | 0.003 | 0.003 | 0.003 | 0.002 | 0.002 | 0.002 | 0.002 | 0.002 | 0.002 | 0.002 |
| Null q *STD* | 0.001 | 0.001 | 0.001 | 0.001 | 0.001 | 0.001 | 0.001 | 0.001 | 0.001 | 0.001 | 0.001 |
| Pseudo-*Z* | 56.035 | 62.069 | 53.849 | 56.745 | 59.171 | 58.568 | 62.189 | **76.389** | 63.611 | 60.320 | 62.596 |
| *P*-value | 0 | 0 | 0 | 0 | 0 | 0 | 0 | 0 | 0 | 0 | 0 |

This table summarizes the results of a permutation-based cluster-quality analysis used to determine the optimal number of clusters for the 659 significant functional connectivity pairs (*n* = 88). For each candidate number of clusters (*k* = 5-15), we computed the mean silhouette value (Quality) of the real data and compared it to a null distribution generated from 100 column-permuted datasets. The highest pseudo-*Z* score identified *k* = 12 as the optimal clustering solution.

**Table S12. Predicting cytokine fold change differences from functional network cluster centroids**

| | *IL-1β* | | | *CXCL8* | | | *TNF-α* | | | *TGF-β1* | | |
|---|---|---|---|---|---|---|---|---|---|---|---|---|
| Cluster | *β* | *t* | *P* | *β* | *t* | *P* | *β* | *t* | *P* | *β* | *t* | *P* |
| 1 | 0.50 | 0.30 | 0.7668 | -5.10 | -1.39 | 0.1714 | 0.62 | 0.52 | 0.6053 | 1.25 | 1.14 | 0.2602 |
| 2 | 0.66 | 0.29 | 0.7733 | -1.72 | -0.34 | 0.7368 | 2.06 | 1.30 | 0.1999 | 1.63 | 1.10 | 0.2773 |
| 3 | -2.09 | -1.18 | 0.2451 | -3.00 | -0.75 | 0.4565 | 1.20 | 0.95 | 0.3470 | 0.81 | 0.69 | 0.4964 |
| 4 | -0.30 | -0.14 | 0.8924 | -4.55 | -0.94 | 0.3507 | 1.83 | 1.20 | 0.2368 | 2.06 | 1.45 | 0.1524 |
| 5 | -2.78 | -1.30 | 0.2001 | -3.85 | -0.80 | 0.4268 | -0.58 | -0.38 | 0.7050 | -0.30 | -0.21 | 0.8374 |
| 6 | 0.40 | 0.21 | 0.8371 | -4.92 | -1.15 | 0.2573 | -0.17 | -0.12 | 0.9042 | 0.93 | 0.73 | 0.4698 |
| 7 | 1.86 | 0.81 | 0.4231 | 14.43 | 3.05 | 0.0036 ** | -2.53 | -1.59 | 0.1192 | -2.35 | -1.58 | 0.1202 |
| 8 | 1.74 | 0.81 | 0.4233 | -1.36 | -0.28 | 0.7790 | 0.14 | 0.09 | 0.9271 | 1.17 | 0.83 | 0.4124 |
| 9 | 1.23 | 0.65 | 0.5193 | 3.40 | 0.81 | 0.4237 | -0.42 | -0.31 | 0.7584 | -0.22 | -0.17 | 0.8645 |
| 10 | 2.99 | 1.41 | 0.1636 | -4.12 | -0.87 | 0.3904 | -0.02 | -0.02 | 0.9870 | 1.06 | 0.75 | 0.4568 |
| 11 | 3.56 | 1.62 | 0.1105 | -2.48 | -0.50 | 0.6213 | 3.09 | 2.03 | 0.0477 * | 1.72 | 1.18 | 0.2435 |
| 12 | -0.47 | -0.26 | 0.7949 | -0.23 | -0.06 | 0.9550 | -0.77 | -0.61 | 0.5427 | -1.11 | -0.95 | 0.3467 |

To test whether cytokine fold differences (ΔFC; VT−VC) could be predicted from functional network cluster centroids, we conducted linear mixed-effect modeling for each cytokine ($n$ = 55). The model formula was: ΔFC ~ Cluster centroid + Condition Order + Sex + Age. We found the centroid of cluster 7 significantly predicted Δfold change of CXCL8 ($P = 0.0036$, Bonferroni-corrected across 12 clusters; **Fig. 5**). *$P < .05$. **$P < .01$.
